# Misoprostol Treatment Prevents Hypoxia-Induced Cardiac Dysfunction Through a 14-3-3 and PKA regulatory motif on Bnip3

**DOI:** 10.1101/2020.10.09.333666

**Authors:** Matthew D. Martens, Nivedita Seshadri, Lucas Nguyen, Donald Chapman, Elizabeth S. Henson, Bo Xiang, Landon Falk, Arielys Mendoza, Sunil Rattan, Spencer B. Gibson, Richard Keijzer, Ayesha Saleem, Grant M. Hatch, Christine A. Doucette, Jason M. Karch, Vernon W. Dolinsky, Ian M. Dixon, Adrian R. West, Christof Rampitsch, Joseph W. Gordon

## Abstract

Systemic hypoxia is a common element in most perinatal emergencies and is a known driver of Bnip3 expression in the neonatal heart. Bnip3 plays a prominent role in the evolution of necrotic cell death, disrupting ER calcium homeostasis and initiating mitochondrial permeability transition (MPT). Emerging evidence suggests a cardioprotective role for the prostaglandin E1 analogue misoprostol during periods of hypoxia, but the mechanisms for this protection are not completely understood. Using a combination of mouse and cell models, we tested if misoprostol is cardioprotective during neonatal hypoxic injury by altering Bnip3 function. Here we report that hypoxia elicits mitochondrial-fragmentation, MPT, reduced ejection fraction, and evidence of necroinflammation, which were abrogated with misoprostol treatment or Bnip3 knockout. Through molecular studies we show that misoprostol leads to PKA-dependent Bnip3 phosphorylation at threonine-181, and subsequent redistribution of Bnip3 from mitochondrial Opa1 and the ER through an interaction with 14-3-3 proteins. Taken together, our results demonstrate a role for Bnip3 phosphorylation in the regulation of cardiomyocyte contractile/metabolic dysfunction, and necroinflammation. Furthermore, we identify a potential pharmacological mechanism to prevent neonatal hypoxic injury.

## 1. Introduction

Systemic hypoxia is a major complication associated with nearly all perinatal emergencies, including placental abnormalities, preterm birth, abnormal and impaired lung development, and cyanotic forms of congenital heart disease. Furthermore, according to the United Nations Inter-agency Group for Child Mortality Estimation, preterm birth, intrapartum-related asphyxia, and birth defects are the leading cause neonatal mortality ^1^. Systemic hypoxia is known to activate pathological hypoxic signaling across most neonatal tissues ^2–4^. Moreover, stressors linked to hypoxic injury have been shown to alter neonatal cardiac metabolism, resulting in diminished contractile performance and compromised tissue perfusion, further compounding neuro-cognitive and end-organ complications ^5^. A lack of oxygen at the level of the cardiomyocyte results in the accumulation and activation of transcription factors belonging to the hypoxia-inducible factor-alpha (HIF) family ^6, 7^. This inducible pathway is conserved throughout life, functioning to drive the expression of a number of genes, and promoting cardiomyocyte glycolysis and diminishing mitochondrial respiration when oxygen tension is low ^6–8^.

Bcl-2-like 19 kDa-interacting protein 3 (Bnip3) is one such hypoxia-inducible gene and is also a member of the pro-apoptotic BH3-only subfamily of the Bcl-2 family of proteins ^9–13^. Bnip3 is an atypical member of this subfamily, relying on a C-terminal transmembrane (TM) domain for its pro-death functions ^14^. Previous studies indicate that through this functionally important domain, Bnip3 inserts through the outer mitochondrial membrane, interacting with optic atrophy-1 (Opa1), driving mitochondrial bioenergetic collapse, fission, cytochrome c release and apoptosis ^14–16^. Additionally, Bnip3 can also localize to the endoplasmic reticulum (ER), interrupting Bcl-2-induced inhibition of inositol trisphosphate receptor (IP_3_R) calcium leak, resulting in ER calcium depletion and mitochondrial matrix calcium accumulation, mediated though voltage-dependent anion channel (VDAC) and the mitochondrial calcium uniporter (MCU) ^14, 17–20^. Accumulating evidence suggests that elevated matrix calcium is an important trigger for mitochondrial permeability transition (MPT), a phenomena that is required for the induction of necrotic cell death and evolution of ischemic injury ^21–28^. Key to both of these processes is Bnip3’s ability to drive mitochondrial bioenergetic collapse by affecting complexes of the inner mitochondrial membrane’s electron transport chain, and ultimately ATP production ^29^. Taken together, these observations have long made Bnip3 an attractive therapeutic target in the heart, but this has been met with limited success.

Prostaglandin signaling has pleiotropic effects on the cardiovascular system. Although generally regarded as pro-inflammatory, prostaglandin E1 (PGE1) has also been associated with the resolution of inflammation, T cell inhibition, and wound healing ^30–33^. Interestingly, previous work has demonstrated that prostaglandin signalling through the EP4 receptor, a G-protein coupled receptor classically associated with enhanced protein kinase A (PKA) activity, improves cardiac function in mice following infarction ^34^. Furthermore, overexpression of the EP3 receptor, known to reduce PKA activity, is deleterious in the murine heart ^35^. Recent work from our group built on this, demonstrating that misoprostol, a PGE1 analogue, activates PKA signaling and is cytoprotective in an *in vitro* model of chronic hypoxia ^10^. Moreover, loss of Bnip3 activity through RNA interference or genetic ablation is protective in the heart and cultured myocytes, respectively following prolonged periods of low oxygen tension ^36, 37^. Previous work in HEK 293 and human carcinoma cell culture models has further concluded that phosphorylation of Bnip3 within its TM domain can inhibit apoptosis; however, the upstream signalling pathways that regulate post-translational modification of Bnip3 have not been previously described ^15^. In addition, it is not currently known if Bnip3 phosphorylation can be pharmacologically targeted to modulate cardiomyocyte permeability transition and *in vivo* heart function during a hypoxic episode ^15^. Based on these previous studies, we examined if prostaglandin signalling though misoprostol treatment is sufficient to alter Bnip3 phosphorylation in order to prevent mitochondrial perturbations and contractile dysfunction in the neonatal heart.

In this report we provide novel evidence that misoprostol inhibits hypoxia-induced neonatal contractile dysfunction resulting from cardiomyocyte respiratory collapse and a necroinflammatory phenotype. We further show that this is a result of inhibiting Bnip3-induced transfer of calcium from the ER to the mitochondria, which prevents mitochondrial dysfunction, ATP depletion, MPT and necrotic cell death. Mechanistically, we provide evidence that this process is regulated through EP4 receptor and downstream PKA signalling, by elucidating a novel phosphorylation site on mouse Bnip3 at threonine-181. We further delineate a role for the 14-3-3 family of molecular chaperones in this pathway, demonstrating that the interaction between 14-3-3β and Bnip3 is increased by misoprostol treatment, facilitating Bnip3 trafficking from the ER and mitochondria. Given the diverse roles of Bnip3 in hypoxic pathologies and cancer, this mechanism may have broader implications to human disease.

## 2. Materials and Methods

### 2.1. In Vivo Neonatal Hypoxia Model and Adult Coronary Ligation Model

All procedures in this study were approved by the Animal Care Committee of the University of Manitoba, which adheres to the principles for biomedical research involving animals developed by the Canadian Council on Animal Care (CCAC). Litters of wild-type and/or Bnip3-null (embryonic deletion described previously ^37^) C57BL/6 mouse pups and their dams were placed in a hypoxia chamber with 10% O_2_ (±1%) on postnatal day (PND) 3 for a period of 7 consecutive days. Control litters were left in normoxic conditions at 21% O_2_. Animals received 10 μg/kg misoprostol or saline control, administered through subcutaneous injection daily from PND3-10. At PND10 animals were euthanized and perfused with saline for tissue collection. This hypoxia protocol has been previously shown to induce cognitive impairment consistent with hypoxic-ischemic encephalopathy ^38, 39^. In the *in vivo* rodent model of myocardial infarction, the left coronary artery of Sprague Dawley rats was ligated approximately 2 mm from its origin, while sham operated rats serve as control ^40, 41^. Following recovery for 4 or 8 weeks, animals are anesthetized, the heart excised, and the left anterior descending territory dissected for scar tissue and viable border-zone myocardium.

### 2.2. In Vivo Assessment of Cardiac Function

Transthoracic echocardiography was performed on mildly anesthetized rats (sedated with 3% isoflurane in oxygen at 1 L/min and maintained at 1-1.5% isoflurane in oxygen at 1 L/min) at PND10 using a Vevo 2100 High-Resolution Imaging System equipped with a 30-MHz transducer (RMV-716; VisualSonics, Toronto) as described previously ^42^.

### 2.3 Cell Culture and Transfections

Rat primary ventricular neonatal cardiomyocytes (PVNC) were isolated from 1-2-day old pups using the Pierce Primary Cardiomyocyte Isolation Kit (#88281), which includes a cardiomyocyte growth supplement to reduce fibroblast contamination. H9c2 cells were maintained in Dulbecco’s modified Eagle’s medium (DMEM; Hyclone), containing penicillin, streptomycin, and 10% fetal bovine serum (Hyclone). Media was supplemented with MEM Non-Essential Amino Acids Solution (Gibco) for MEFs. Cells were incubated at 37°C and 5% CO2. Human induced pluripotent stem cell-derived cardiomyocytes (H-iPSC-CMs) were obtained from Cellular Dynamics (iCell Cardiomyocytes #01434). iCell Cardiomyocytes were cultured in maintenance medium as per the manufacturer’s protocol and differentiated for 72 hours. Cell lines were transfected using JetPrime Polyplus reagent, as per the manufacturer’s protocol. For misoprostol treatments, 10 mM misoprostol (Sigma) in phosphate buffered saline (PBS; Hyclone) was diluted to 10 μM directly in the media and applied to cells for 24 hours. To achieve hypoxia, cells were held in a Biospherix incubator sub-chamber with 1% O_2_ (±1%), 5% CO_2_, balanced with pure N_2_ (regulated by a Biospherix ProOx C21 sub-chamber controller) at 37 °C for 24 hours. BvO2, L161-982, L798-106, and H89 dihydrochloride (H89) were purchased from Sigma. RNAi experiments targeting Bnip3, including the generation of si-Bnip3 were described previously ^10^.

### 2.4 Plasmids

Mito-Emerald (mEmerald-Mito-7) was a gift from Michael Davidson (Addgene #54160) ^43^. The endoplasmic reticulum (CMV-ER-LAR-GECO1), and mitochondrial (CMV-mito-CAR-GECO1) targeted calcium biosensors were gifts from Robert Campbell (Addgene #61244, and #46022) ^44^. CMV-dsRed was a gift from John C. McDermott. The FRET-based ATP biosensor (ATeam1.03-nD/nA/pcDNA3) was a gift from Takeharu Nagai (Addgene plasmid #51958) ^45^. The dimerization-dependent PKA biosensor (pPHT-PKA) was a gift from Anne Marie Quinn (Addgene #60936) ^46^. pcDNA3-HA-14-3-3 beta (14-3-3β) was a gift from Michael Yaffe (Addgene #13270). pclbw-opa1(isoform 1)-myc (myc-Opa1) was a gift from David Chan (Addgene plasmid # 62845) ^47^. The generation of mouse myc-Bnip3 (Addgene #100796) was described previously ^48^. The phospho-neutral mouse myc-Bnip3-T181A was generated by PCR using the New England Biolabs Q5 Site-Directed Mutagenesis Kit and primers Forward: 5’-CTAGTCTAGA ATGTCGCAGAGCGGGGAGGAGAAC-3’and Reverse: 5’-GATCGGATCCTCAGAAGGTGCTAGTGGAAGTtgcCAG-3’.

### 2.5 Fluorescent Staining, Live Cell Imaging and Immunofluorescence

MitoView Green, TMRM, Calcein-AM, ethidium homodimer-1, and Hoechst 33342 were purchased from Biotium. MitoSox was purchased from Life Technologies. MPTP imaging was described previously ^24^. Dye based calcium imaging was done with Rhod-2AM (Invitrogen, R1245MP) as per manufacturer’s protocol (including the production of dihyrdorhod-2 AM). Immunofluorescence with HMBG1 (CST # 3935), Bnip3 (CST # 3769 and Ab-196706), 14-3-3β (sc-25276), and Opa1 (sc-393296) antibodies were performed in conjunction with fluorescent secondary antibodies conjugated to Alexa Fluor 466 or 647 (Jackson). All epifluorescent imaging experiments were done on a Zeiss Axiovert 200 inverted microscope fitted with a Calibri 7 LED Light Source (Zeiss) and Axiocam 702 mono camera (Zeiss) in combination with Zen 2.3 Pro imaging software. Confocal imaging was done on a Zeiss LSM700 Spectral Confocal Microscope in combination with Zen Black, which was also used for colocalization analysis, while FRET imaging was done using a Cytation 5 Cell Imaging Multi-Mode Reader. Quantification, scale bars, and processing including background subtraction, was done on Fiji (ImageJ) software.

### 2.6. Transmission Electron Microscopy (TEM)

TEM imaging was performed according to a protocol described previously ^49^. Briefly, PND10 hearts were fixed (3% glutaraldehyde in PBS, pH 7.4) for 3 hours at room temperature. Hearts were treated with a post-fixation step using 1% osmium tetroxide in phosphate buffer for 2 hours at room temperature, followed by an alcohol dehydration series before embedding in Epon. TEM was performed with a Philips CM10, at 80 kV, on ultra-thin sections (100 nm on 200 mesh grids). Hearts were stained with uranyl acetate and counterstained with lead citrate.

### 2.7. Immunoblotting

Protein isolation and quantification was performed as described previously ^10^. Extracts were resolved via SDS-PAGE and later transferred to a PVDF membrane using an overnight transfer protocol. Immunoblotting was carried out using primary antibodies in 5% powdered milk or BSA (as per the manufacturer’s instructions) dissolved in TBST. Horseradish peroxidase-conjugated secondary antibodies (Jackson ImmunoResearch Laboratories; 1:5000) were used in combination with enhanced chemiluminescence (ECL) to visualize bands. The following antibodies were used: HMGB1 (CST # 3935), Rodent-specific Bnip3 (CST # 3769), Myc-Tag (CST # 2272), HA-Tag (CST # 3724), AIF (CST # 5318), MEK1/2 (CST # 8727), SERCA (Sigma MA3-919), Actin (sc-1616), and Tubulin (CST #86298). For detection of phosphorylation of Bnip3 at threonine-181, a custom rabbit polyclonal antibody was generated by Abgent using the following peptide sequence: AIGLGIYIGRRLp(T)TSTSTF.

### 2.6. Quantitative PCR

Total RNA was extracted from pulverized frozen tissue or from cultured cells by TRIzol method. cDNA was generated using SuperScript IV VILO Master Mix with ezDNase (Thermo #11766050) and q-RT-PCR performed using TaqMan Fast Advanced master mix (Thermo #4444965) on a CFX384 Real-Time PCR Instrument. The primers used were provided through ThermoFisher custom plating arrays (see Supplement 1 and 2 for assay list).

### 2.7. Cardiac and Cellular Lactate, ATP and Extracellular Acidification

Cardiac lactate was quantified using the bioluminescent Lactate-Glo™ Assay (Promega #J5021) in deproteinized PND10 heart samples, as per the manufacturer’s protocol. Luminescence was detected and quantified using a Fluostar Optima microplate reader (BMG Labtech, Ortenberg, Germany). Cardiac and H9c2 ATP content was determined using a the Adenosine 5′-triphosphate (ATP) Bioluminescent Assay Kit (Sigma #FLAA-1KT), and normalized to DNA content as described previously ^50^. Extracellular acidification and oxygen consumption was determined on a Seahorse XF-24 Extracellular Flux Analyzer in combination with Seahorse Mitochondrial Stress Test with drug concentrations as follows: 1 uM Oligomycin, 2 μ M FCCP and 1 μM rotanone/antimycin A (Agilent Seahorse #103015-100). Calculated oxygen consumption rates were determined as per the manufacturer’s instructions (Mitochondrial Stress Test; Seahorse).

### 2.8. Mitochondrial Swelling and CRC Assays

Mitochondrial swelling and calcium retention capacity (CRC) assays were performed using a cuvette–based fluorometric system (Horiba Scientific) which allows for the simultaneous detection of both fluorescence and absorbance, as described previously ^51^. Hearts were minced in mitochondrial isolation buffer and homogenized using a 2 mL Teflon-glass homogenizer. Mitochondria were enriched by differential centrifugation at 4°C. The mitochondrial isolation buffer consisted of 225 mM mannitol, 75 mM sucrose, 5 mM HEPES, and 1 mM EGTA, pH 7.4. Within the cuvette, 2 mg of mitochondria were incubated with 250 nM Calcium Green-5N (Invitrogen), 7 mM pyruvate (Sigma-Aldrich), and 1 mM malate (Sigma-Aldrich) in 1 ml of hypotonic KCl buffer (125 mM KCl, 20 mM HEPES, 2 mM KH2PO4, 40 μM EGTA, pH 7.2). In some experiments, mitochondria were incubated with 10 μM misoprostol for 5 minutes prior to the start of the assay. Mitochondria were then pulsed with sequential additions of CaCl2 (20 μM) over specific increments of time until mitochondrial swelling occurred.

### 2.9. Phospho Peptide Mapping

Synthetic peptides (GeneScript) were resuspended at a concentration of 1 mg/ml. These peptides were used as the substrate in a PKA kinase assay kit (New England Biolabs, #P6000S) according to the manufacturer’s instructions, with the exception that [32P]-ATP was replaced with fresh molecular biology grade ATP. The Kemptide substrate (Enzo Life Sciences; #P-107; LRRASLG) was used as a positive control in each assay. Before mass spectrometry analysis, kinase assays were prepared using C18 ZipTips (EMD Millipore, Etobicoke, ON, Canada). Samples in 50% acetonitrile and 0.1% formic acid were introduced into a linear ion-trap mass spectrometer (LTQ XL: ThermoFisher, San Jose, CA, USA) via static nanoflow, using a glass capillary emitter (PicoTip: New Objective, Woburn, MA, USA), as described previously ^52^.

### 2.10. Statistics

Data are presented as mean ± standard error (S.E.M.) from 3 independent cell culture experiments. Differences between groups in imaging experiments with only 2 conditions were analyzed using an unpaired t-test, where (*) indicates P<0.05 compared with control. Experiments with 4 or more conditions were analyzed using a 1-way ANOVA, or 2-way ANOVA where appropriate, with subsequent Tukey test for multiple comparisons, where (*) indicates P<0.05 compared with control, and (**) indicates P<0.05 compared with treatment. All statistical analysis was done using GraphPad Prism software.

## 3. Results

### 3.1. Misoprostol prevents hypoxia-induced contractile and mitochondrial dysfunction in vivo

Given that cardiac dysfunction has been clinically implicated in neonatal hypoxic injury, we evaluated cardiac function in a mouse model of HIE (10% oxygen from PND 3-10)(Fig. 1 A)^10, 53^. Transthoracic echocardiography revealed signs of significant contractile dysfunction in hypoxia exposed neonatal animals, which included reductions in both fractional shortening (FS) and ejection fraction (EF), alongside considerable alterations in the ability for the left ventricle to fill properly in between beats (E’/A’) (Fig. 1 B-D). However, when hypoxic animals were concurrently treated with misoprostol, contractility and filling of the heart returned to normoxic control levels (Fig. 1 B-D).

**Figure 1.**
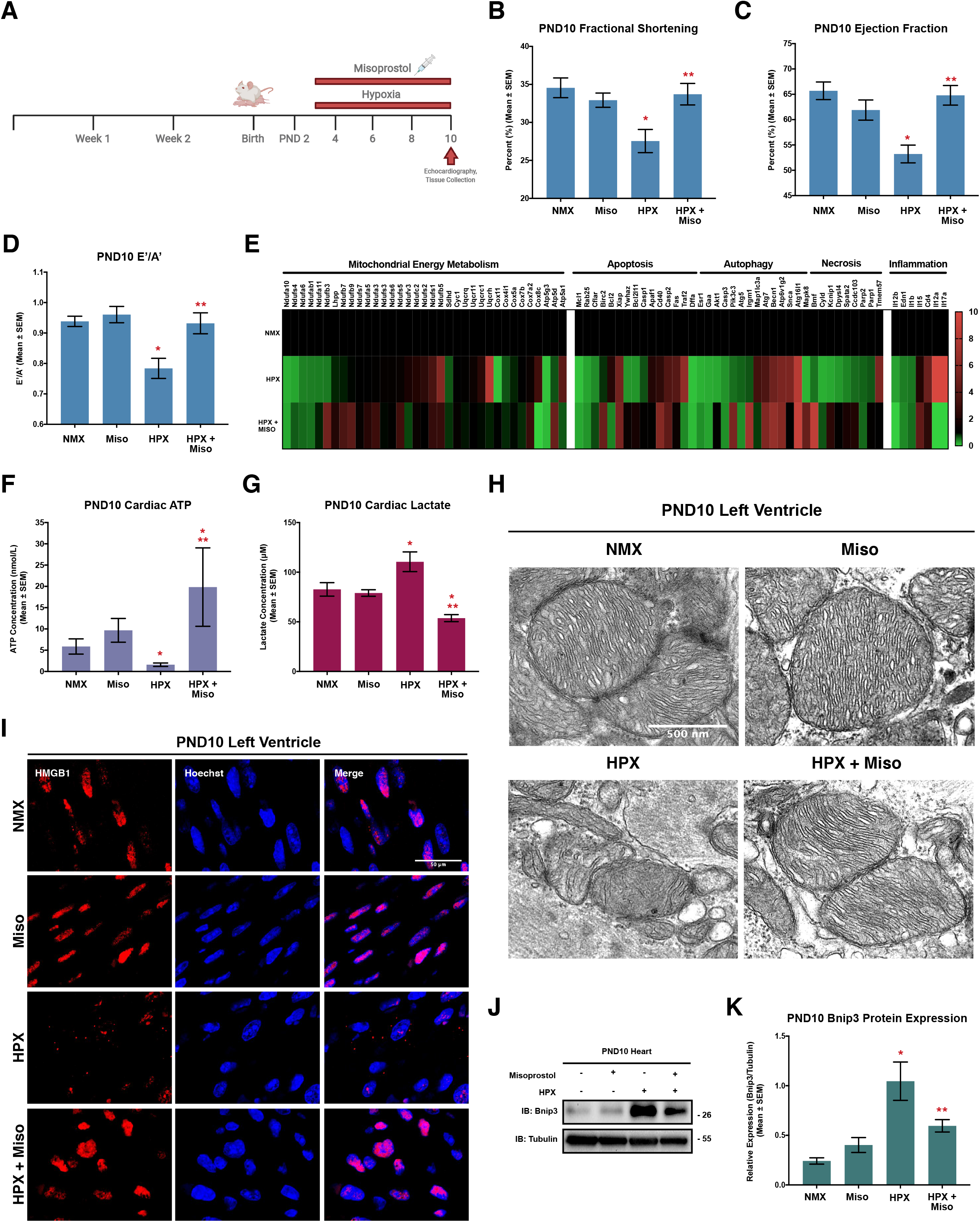
Misoprostol prevents hypoxia-induced contractile and mitochondrial dysfunction in vivo. (**A**) Schematic of the mouse model of neonatal hypoxia, where mice are exposed to hypoxia (10% O_2_) ± 10 μg/kg misoprostol daily from PND3-10. **(B)** Fractional shortening, **(C)** Ejection fraction, and **(D)** E’/A’ ratio, for 4-6 post-natal day (PND10) mice treated as in (A), as determined by transthoracic echocardiography. **(E)** PCR-based array performed on RNA isolated from PND10 mouse ventricles (n=3 animals per group) treated as in (A), where green indicates a downregulation of expression (<1), and red indicates an upregulation of expression (>1), relative to the normoxic control (1). **(F)** Measurement of ATP content in PND10 mouse ventricles (n=6-8 animals per condition) treated as in (A). **(G)** Measurement of cardiac lactate content in the PND10 mouse ventricle (n=6-8 animals per condition) treated as in (A). **(H)** PND10 hearts treated as in (A) and imaged via transmission electron microscopy. Images showing mitochondrial morphology. **(I)** PND10 hearts treated as in (A) and stained with DAPI (Blue) and probed for high mobility group box 1 (HMGB1, red). Hearts were imaged via confocal microscopy. **(J)** Representative immunoblot of heart protein extracts from post-natal day (PND10) mice treated as in (A). Extracts were immunoblotted as indicated. **(K)** Relative Bnip3 gene expression from the PND10 mouse ventricles of animals (n=6-9 animals per group) treated as in (A). All data are represented as mean ± S.E.M. *\*P*<0.05 compared with control, while *\*\*P*<0.05 compared with hypoxia treatment, determined by 1-way ANOVA or 2-way ANOVA where appropriate.

In an effort to understand the underlying mechanism of this dysfunction, we ran targeted PCR arrays to assess the expression of genes associated with mitochondrial energy metabolism, various cell death pathways, and genes expressing inflammatory cytokines (Fig. 1 E). Through this approach we found that there was a change in the abundance of mRNAs for proteins associated with the mitochondrial electron transport chain (ETC); however, very few of these genes were returned to control levels with misoprostol treatment. In particular there was a downregulation of mRNAs associated with complex-1 (NADH ubiquinone oxidoreductase) (Fig.1 E), a phenomena which has previously been linked to mitochondrial dysfunction, fragmentation and bioenergetic collapse ^29^. Additionally, we observed that mRNAs associated with ATP synthase, the terminal step of electron transport and main beneficiary of a well-developed proton motive force, were also less abundant in the hypoxic neonatal heart, relative to normoxic control animals (Fig. 1 E). When we examined cell death pathways, we found that a number of regulators of apoptosis and necrosis were reduced in abundance, including induced myeloid leukemia cell differentiation protein (Mcl-1), Ras-Related Protein Rab-25 (Rab25), CASP8 and FADD Like Apoptosis Regulator (Cflar) (Fig. 1E). At the same time hypoxia increased the abundance of mRNAs that drive extrinsic cell death and inflammation like, Tumor Necrosis Factor Receptor Superfamily Member 5 (CD40), Fas Cell Surface Death Receptor (Fas), TNF Receptor Associated Factor 2 (Traf2), and the DNA Fragmentation Factor Subunit Alpha (Dffa). Interestingly, interleukins (IL) 17a and 12a, involved in innate-and T cell-mediated inflammation, were increased during hypoxic injury and reduced below detectable levels by concurrent misoprostol treatment (Fig. 1 E).

Next, we determined if the hypoxic neonatal heart demonstrated signs of mitochondrial dysfunction. To do this, we assessed cardiac ATP content, and observed that after 7 days of hypoxia exposure, there was a significant reduction in ATP in the neonatal heart (Fig. 1 F). Importantly, when we combined hypoxia with misoprostol treatment, ATP content was elevated significantly beyond that observed in the normoxic control animals (Fig. 1 F). We next assessed lactate content in the neonatal heart using a bioluminescent assay, and observed that there was a 33% increase in lactate concentration in hypoxic animals, when compared to controls (Fig. 1 G). However, daily misoprostol treatment during exposure to hypoxia was sufficient to significantly reduce lactate accumulation in the hypoxic mouse heart (Fig. 1 G). We then used transmission electron microscopy (TEM) to visualize mitochondrial morphology. This analysis revealed that hypoxia both reduced mitochondrial size and altered their structure when compared to normoxic controls; however, these changes were prevented when hypoxic animals were treated with misoprostol (Fig.1 H).

Building on the biochemical indicators of bioenergetic collapse and the increased abundance of mRNAs for proteins involved necrosis and inflammation, we examined the subcellular distribution of high mobility group box 1 (HMGB1), a nuclear protein that is released into the cytosol and interstitium during necrosis where it acts as an alarmin or damage-associated molecular pattern (ie. DAMP) to engage an innate immune/inflammatory response. Using confocal immunofluorescence, hypoxia exposed left ventricles demonstrated a marked decrease in HMGB1 nuclear localization; however, when we combined hypoxia with daily misoprostol drug treatments, we observed that HMGB1 remained in the nucleus (Fig. 1 I). Given the central role of Bnip3 in hypoxia-induced mitochondrial permeability transition, bioenergetic collapse, and necrotic cell death, we also assessed its expression by Western blot analysis, and observed that Bnip3 expression is significantly increased in the hypoxia-exposed neonatal heart, and partially reduced by misoprostol treatment (Fig. 1 J, K).

Together these results indicate that the hypoxia exposed neonatal heart may be undergoing bioenergetic collapse leading to a necroinflammatory phenotype resulting in significant alterations in contractile function. Interestingly, treatment with misoprostol at least transiently maintains mitochondrial morphology and function, despite hypoxia-induced changes in mitochondrial gene expression. Moreover, misoprostol can prevent the necroinflammatory response elicited by hypoxia, with an overall preservation in cardiac function. Finally, although previous reports have demonstrated that misoprostol can repress hypoxia-induced Bnip3 expression in other tissues, misoprostol treatment only partially reduced Bnip3 in the neonatal heart, suggesting that additional pharmacodynamic mechanisms operate in this organ.

### 3.2. Misoprostol prevents hypoxia-induced mitochondrial dysfunction in rodent and human cardiomyocytes

In order to investigate if misoprostol could modulate hypoxia-induced mitochondrial dysfunction and necrosis at the cellular level, we employed cultured primary ventricular neonatal cardiomyocytes (PVNCs), isolated from PND 1–2 rat pups. When exposed to hypoxia for 24 hours we observed a significant increase in mitochondrial fragmentation when compared to normoxic control cells (Fig. 2 A, B), consistent with our TEM data, and commonly associated with mitochondrial dysfunction and complex-1 deficiency. However, when PVNCs were concurrently exposed to hypoxia and treated with misoprostol, fragmentation was absent, and mitochondria returned to their normal branching and networked appearance (Fig. 2 A, B).

**Figure 2.**
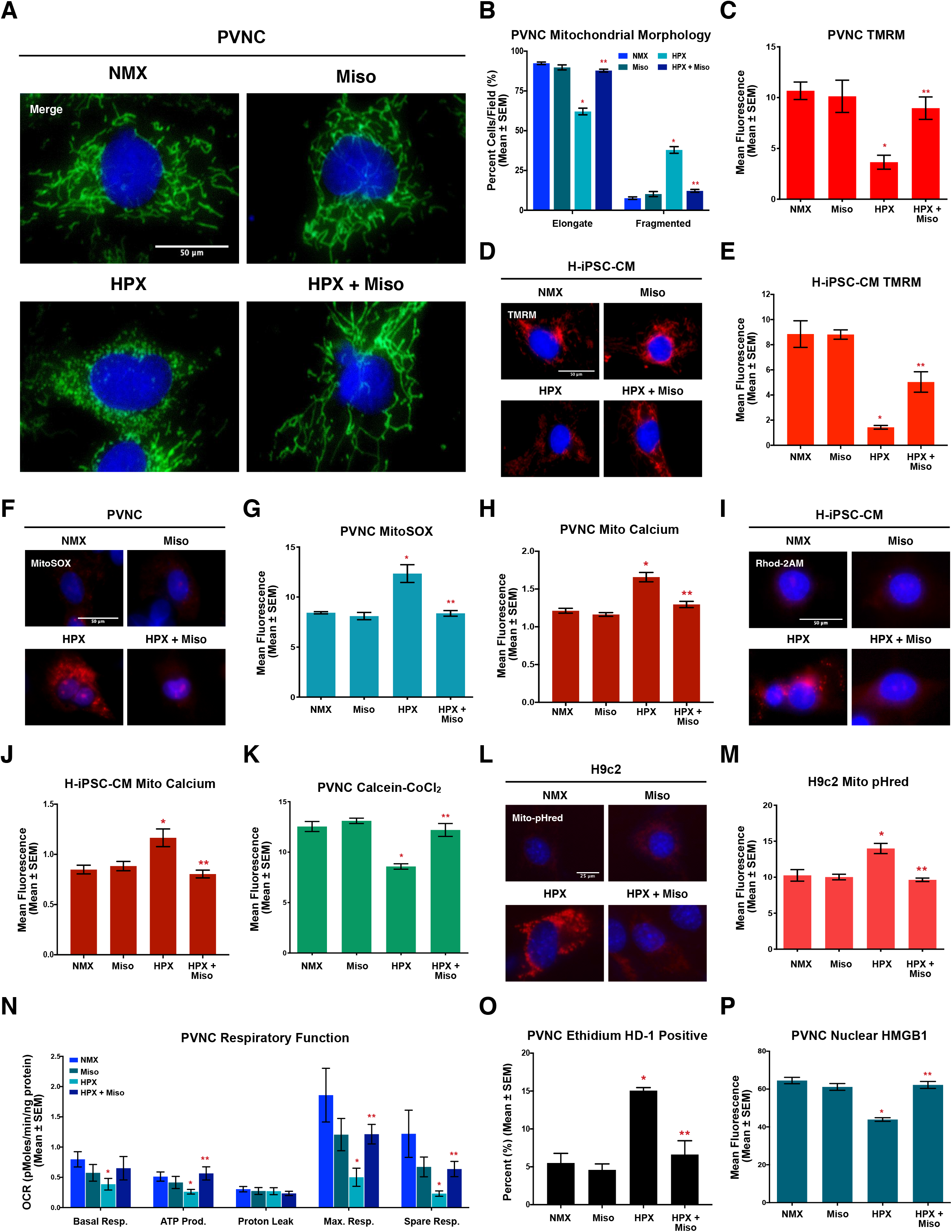
Misoprostol prevents hypoxia-induced mitochondrial dysfunction *in vitro*. **(A)** Primary ventricular neonatal cardiomyocytes (PVNCs) treated with 10 μM misoprostol (Miso) ± 1% O_2_ (HPX) for 24 hours. MitoView Green was included in all conditions to show mitochondrial morphology. Cells were stained with hoechst (blue) and imaged by standard fluorescence microscopy. **(B)** Quantification of cells in (A), where the number of cells with elongated and fragmented mitochondria are expressed as a percentage of all transfected cells in 30 random fields, across 3 independent experiments. **(C)** Quantification of PVNC’s treated as in (A). Cells were stained with TMRM (red) and hoechst (blue) and imaged by standard fluorescence microscopy. Red fluorescent signal was normalized to cell area and quantified in 30 random fields, across 3 independent experiments. **(D)** Human induced pluripotent stem cell-derived cardiomyocytes (H-IPSC-CMs) treated as in (A). Cells were stained with TMRM (red) and hoechst (blue) and imaged by standard fluorescence microscopy. **(E)** Quantification of cells in (D), as in (C). **(F)** PVNC’s treated as in (A). Cells were stained with MitoSOX (Red) and hoechst (blue) and imaged by standard fluorescence microscopy. **(G)** Quantification of cells in (F), as in (C). **(H)** Quantification of PVNC’s treated as in (A). Cells were stained with dihyrorhod-2AM (Red) and hoechst (blue) and imaged by standard fluorescence microscopy and quantified as in (C). **(I)** H-IPSC-CMs treated as in (A). Cells were stained with dihyrorhod-2AM (Red) and hoechst (blue) and imaged by standard fluorescence microscopy. **(J)** Quantification of cells in (I), as in (C). **(K)** Quantification of PVNC’s treated as in (A). Cells were stained with hoechst (blue) and calcein-AM quenched by cobalt chloride (CoCl_2_, 5 μM) to assess permeability transition, where green fluorescent signal was normalized to cell area and quantified in 30 random fields, across 3 independent experiments. **(L)** H9c2 cells treated as in (A), GW-1-Mito-pHred (red) was included in all conditions to visualize mitophagy. Cells were stained with hoechst (blue) and imaged by standard fluorescence microscopy. **(M)** Quantification of cells in (L), where red fluorescent signal was normalized to cell area and quantified in 15 random fields, across 3 independent experiments. **(N)** Calculated oxygen consumption rates (OCR) determined by Seahorse Xf-24 analysis to evaluate mitochondrial function. **(O)** Quantification of PVNC’s treated as in (A). Live cells were stained with calcein-AM (green), and necrotic cells were stained with ethidium homodimer-1 (red). Percent (%) dead was calculated across 30 random fields, across 3 independent experiments. **(P)** Quantification of PVNC’s treated as in (A). Cells were fixed, stained with hoechst (blue), and immunofluorescence was performed using a HMGB1 primary antibody (green). Cells were then imaged by standard fluorescence microscopy. Green fluorescent signal was then normalized to nuclear area and quantified in 30 random fields, across 3 independent experiments. All data are represented as mean ± S.E.M. *\*P*<0.05 compared with control, while *\*\*P*<0.05 compared with hypoxia treatment, determined by 1-way ANOVA or 2-way ANOVA where appropriate.

Due to the previously reported association between fragmentation and mitochondrial dysfunction, we next performed a number of functional assays to assess the response of PVNC mitochondria to hypoxia. To do this we used TMRM, a cell-permeant red fluorescent dye to assess mitochondrial membrane potential (Δm). We observed that hypoxia significantly reduced Δm normoxic controls, which was restored by misoprostol treatment (Fig. 2 C). We were further interested in determining if this observation translated to hypoxia-exposed human induced pluripotent stem cell (IPSC)-derived cardiomyocytes (H-iPSC-CMs). We observed that hypoxia exposure significantly reduced mitochondrial membrane potential in these H-iPSC-CMs, and misoprostol was able to prevent this effect, consistent with the PVNC results (Fig. 2 D, E). We next moved to assess the production of mitochondrial derived superoxide, a common and potent cytotoxic free radical, using MitoSOX staining. We observed that hypoxia markedly increased mitochondrial superoxide production well above that of control levels but this response was completely abrogated in the presence of misoprostol (Fig. 2 F, G).

Next, we investigated how hypoxia alters subcellular calcium dynamics and impacts mitochondrial permeability transition. To do this, we stained cardiomyocytes with a reduced form of Rhod-2AM (dihydrorhod2-AM), which provides specificity for mitochondrial calcium imaging. We observed that 24 hours of hypoxia significantly increased mitochondrial calcium content relative to that of the normoxic controls in both PVNCs and H-iPSC-CMs (Fig. 2 H-J). However, when hypoxic cardiomyocytes were co-treated with misoprostol, the observed hypoxia-dependent increase in mitochondrial calcium was prevented (Fig. 2 H-J). Given the previously published links between mitochondrial calcium accumulation and MPT, we assessed MPT in hypoxic cardiomyocytes using the calcein-CoCl_2_ method. Consistent with the calcium results, hypoxia exposure resulted in a loss of mitochondrial puncta, indicative of permeability transition, while cells that were concurrently treated with hypoxia and misoprostol maintained mitochondrial staining comparable to control levels (Fig. 2 K). Using a mitochondrial-targeted biosensor that fluoresces red in the presence of acidic pHs, like those found in the lysosome, we also assessed the effects of hypoxia on mitophagy in H9c2 cells. In doing this we found that hypoxia significantly increased mitochondrial autophagy; however, this was prevented in the presence of misoprostol (Fig. 2 H, I).

Using extracellular flux analysis, we examined the effects of this mitochondrial dysfunction on oxidative-phosphorylation. As shown in Figure 2N, myocyte hypoxia significantly reduced basal, maximal and spare respiratory capacity, which further resulted in a significant reduction in mitochondrial ATP production. However, consistent with what was seen *in vivo*, the addition of misoprostol during hypoxia abrogated this effect, preventing respiratory collapse in primary neonatal cardiomyocytes (Fig. 2 N).

While these results demonstrate that hypoxia results in mitochondrial dysfunction, we also wanted to determine if the downstream result was ultimately cell death. In order to do this we performed live/dead assays using ethidium homodimer-1 to mark the nuclei of cells that had lost their membrane integrity, a common characteristic of necrotic cell death ^54^. With this approach we observed that hypoxia significantly increased the percentage of red staining nuclei by 173% when compared to normoxic controls; however, myocytes treated with misoprostol displayed levels of cell death similar to that of control cells (Fig. 2 O). To further understand the underlying mechanism of cell death in hypoxia-induced cardiomyocyte pathology we also assessed HMGB1 subcellular localization, an alarmin/DAMP used to assess necrosis. While hypoxic PVNCs demonstrated a significant decrease in nuclear HMGB1 immunofluorescence, the addition of misoprostol during hypoxia restored HMGB1 to the nucleus, resembling the distribution observed in normoxic cells (Fig. 2P). Taken together, these results indicate that misoprostol prevents hypoxia-induced mitochondrial dysfunction, necrotic cell death, and alarmin/DAMP release in neonatal cardiomyocytes.

### 3.3. Misoprostol prevents Bnip3-induced mitochondrial dysfunction and cell death

Given the central role that Bnip3 has previously been shown to play in the evolution of hypoxic injury in the heart, we also wanted to assess its expression in our cellular model ^10, 37^. Using PVNCs we observed that hypoxia exposure increased Bnip3 immunoreactivity (Fig. 3 A, B). Consistent with our previously published results, the addition of misoprostol to PVNCs during hypoxia reduced Bnip3 immunofluorescence, however it still remained elevated relative to the levels observed in the normoxic control cells (Fig. 3 A, B) ^10^.

**Figure 3.**
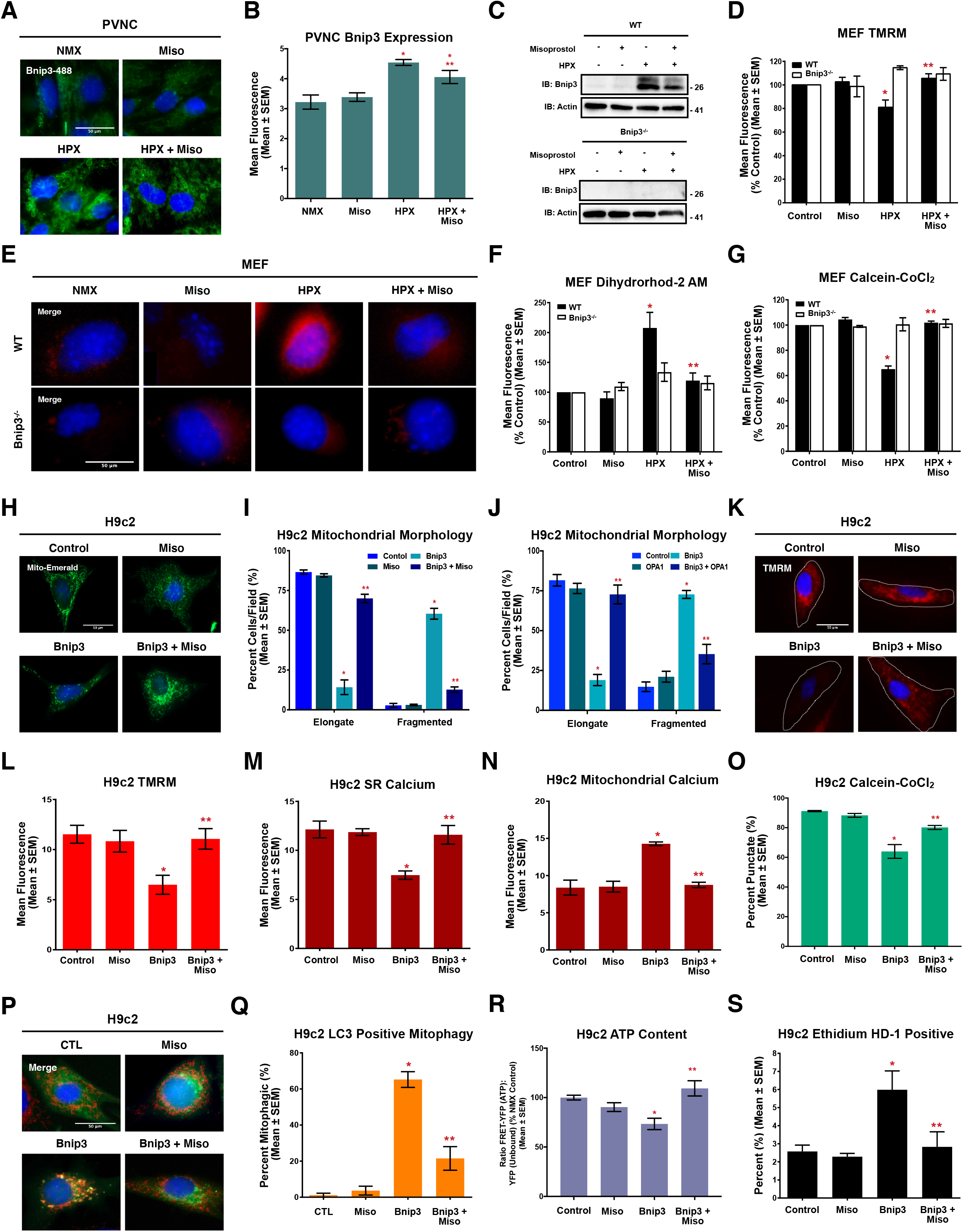
Misoprostol prevents Bnip3-induced mitochondrial dysfunction and cell death. **(A)** PVNC’s treated with 10 μM misoprostol (Miso) ± 1% O_2_ (HPX) for 24 hours. Cells were fixed, stained with hoechst (blue), and immunofluorescence was performed using a Bnip3 primary antibody (green). Cells were then imaged by standard fluorescence microscopy. **(B)** Quantification of cells in (A), where green fluorescent signal was normalized to cell area and quantified in 30 random fields, across 3 independent experiments. **(C)** Immunoblot for Bnip3 in protein extracts from WT and Bnip3^-/-^ MEFs treated as in (A). **(D)** Quantification of WT and Bnip3^-/-^ mouse embryonic fibroblasts (MEFs) treated as in (A). Cells were stained with TMRM (red) and hoechst (blue) and imaged by standard fluorescence microscopy. Red fluorescent signal was normalized to cell area and quantified in 15 random fields, across 3 independent experiments. **(E)** WT and Bnip3^-/-^ MEFs treated as in (A). Cells were stained with hoechst (blue) and dihydrorhod-2AM to stain mitochondrial calcium. **(F)** Quantification of (E) as in (D) in 15 random fields, across 3 independent experiments. **(G)** Quantification of WT and Bnip3^-/-^ MEFs treated as in (A). Cells were stained with hoechst (blue) and calcein-AM quenched by cobalt chloride (CoCl_2_, 5 μM) to assess permeability transition. Green fluorescent signal was normalized to cell area and quantified in 15 random fields, across 3 independent experiments. **(H)** H9c2 cells transfected with pcDNA3 (control) or Myc-Bnip3 and treated with 10 μM misoprostol (Miso) or PBS control for 16 hours. Mito-Emerald (green) was included in all conditions to show transfected cells and mitochondrial morphology. Cells were stained with hoechst (blue) and imaged by standard fluorescence microscopy. **(I)** Quantification of cells in (H), where the number of cells with elongated and fragmented mitochondria are expressed as a percentage of all transfected cells in 30 random fields, across 3 independent experiments. **(J)** Quantification of H9c2 cells transfected with pcDNA3 (control), Myc-Bnip3 and/or myc-Opa1. Mito-Emerald (green) was included in all conditions as in (H) and cells were stained with hoechst (blue) and imaged by standard fluorescence microscopy. Quantification as in (I). **(K)** H9c2 cells treated as in (H). CMV-GFP (outline) was included in all conditions to indicate transfected cells Cells were stained and imaged as in (D). **(L)** Quantification of cells in (K) as in (D). **(M)** Quantification of H9c2 cells treated as in (H). ER-LAR-GECO (red) was included in all conditions to indicate ER calcium content. Cells were stained and imaged as in (H). Quantification was performed as in (D) in 30 random fields, across 3 independent experiments. **(N)** Quantification of H9c2 cells treated as in (H). Mito-CAR-GECO (red) was included in all conditions to indicate mitochondrial calcium content. Cells were stained and imaged as in (D). Quantification was performed as in (D) in 30 random fields, across 3 independent experiments. **(O)** Quantification of H9c2 cells treated as in (H) and CMV-ds.RED was included in all conditions to indicate transfected cells. Cells were stained, imaged and quantified as in (G) across 30 random fields, across 3 independent experiments. **(P)** H9c2 cells treated as in (H). LC3-GFP (green) was included in all conditions to show transfected cells and autophagic puncta. Cells were stained with hoechst (blue) and MitoTracker red (red) and imaged by standard fluorescence microscopy. **(Q)** Quantification of cells in (P), where the number of cells with LC3-GFP and MitoTracker co-localization are expressed as a percentage of all transfected cells in 10 random fields. **(R)** Quantification of H9c2’s treated as in (H). ATeam was used to indicate cytosolic ATP content. Cells were imaged by FRET-based microscopy. FRET-YFP (ATP) signal was divided by the YFP (unbound biosensor) signal in 15 random fields across 3 independent experiments. **(S)** Quantification of H9c2 cells treated as in (H). Live cells were stained with calcein-AM (green), and necrotic cells were stained with ethidium homodimer-1 (red) and are expressed as percent (%) dead in 30 random fields, across 3 independent experiments. All data are represented as mean ± S.E.M. *\*P*<0.05 compared with control, while *\*\*P*<0.05 compared with hypoxia treatment, determined by 1-way ANOVA or 2-way ANOVA where appropriate.

To determine the necessity between Bnip3 function and hypoxia-induced alterations in mitochondrial function, we used fibroblasts isolated from mouse embryos possessing a genetic deletion for Bnip3 (Bnip3^-/-^ MEFs), which were described previously ^12, 37^. Using western blot analysis, we observed that hypoxia increased Bnip3 expression in the wild-type (WT) cells, but not in the Bnip3^-/-^ MEFs (Fig. 3 C). TMRM analysis revealed that in WT MEFs hypoxia significantly reduced Δ m when compared to normoxic control cells, a phenomenon that was absent in the Bnip3^-/-^ MEFs (Fig. 3 D). Importantly, misoprostol also restored mitochondrial membrane potential to control levels in WT MEFs (Fig. 3C, -D). We next assessed mitochondrial calcium accumulation using dihydrorhod2-AM. Hypoxia increased mitochondrial calcium in the WT MEFs, but not in the Bnip3^-/-^ MEFs. Consistent with our PVNC results, misoprostol provided protection against mitochondrial calcium accumulation in the WT cells (Fig. 3 E, F). In addition, Bnip3^-/-^ MEFs were less susceptible to hypoxia-induced MPT compared to WT MEFs, which demonstrated a significant reduction in mitochondrial puncta in response to hypoxia. These effects of hypoxia were similarly prevented by misoprostol (Fig. 3 G).

Given that Bnip3 expression is elevated in our cellular and *in vivo* models of hypoxia, but was not completely repressed by misoprostol treatment, we next determined if misoprostol had a direct effect on Bnip3 function. To do this we used gain-of-function transfection studies in H9c2 cells and assessed markers of mitochondrial dysfunction. H9c2 cells were transfected with Bnip3, or empty vector control, along with mito-Emerald to visualize mitochondrial morphology. As shown in Figure 3 H and I, Bnip3 expression alone significantly altered mitochondrial morphology, resulting in a more fragmented appearance overall. Importantly, when Bnip3-expressing cells were treated with misoprostol this effect was lost and mitochondria retained a branching and networked appearance (Fig. 3 H, I). Given the role of Opa1 in the regulation of mitochondrial morphology, and its documented interaction with Bnip3, we also tested if overexpression of Opa1 rescues Bnip3-induced mitochondrial fragmentation. In doing this we observed that Bnip3 expression consistently resulted in mitochondrial fragmentation, but this effect was lost with the co-expression of Opa1. Using TMRM we additionally observed that Bnip3 expression induced mitochondrial depolarization, which resulted in a 43% reduction in Δm when compared to control (Fig. 3 K, L). However, Bnip3 was unable to reduce the mitochondrial membrane potential in the presence of misoprostol (Fig. 3 K, L).

We next investigated the underlying mechanism of Bnip3-induced mitochondrial dysfunction, focusing on the role of subcellular calcium. To do this we employed organelle-targeted genetically encoded calcium biosensors (GECOs) that fluoresce red in the presence of calcium. When we expressed the ER-targeted calcium biosensors (ER-LAR-GECO) in H9c2 cells, we observed that Bnip3 expression significantly reduced ER calcium stores, when compared to control (Fig. 3 M). Concurrently, data generated using the mitochondrial targeted calcium indicator (Mito-CAR-GECO) demonstrated a Bnip3-dependent increase in mitochondrial calcium (Fig. 3 N). Together this data suggests that Bnip3-induces a shift of calcium from the ER to the mitochondria, consistent with previous observations in a neuronal cell line ^17^. We further demonstrated that this shift in calcium was completely prevented by misoprostol (Fig. 3 M, N). We also evaluated if Bnip3-induced mitochondrial calcium accumulation was impacting MPT. Using calcein staining with cobalt chloride, we observed that Bnip3 expression significantly reduced the number of cells with mitochondrial puncta by nearly 30%, indicating that MPT was actively occurring in these cells (Fig.3 O). In addition, we confirmed that misoprostol prevented Bnip3-induced MPT in H9c2 cells (Fig. 3 O). Additionally, we assessed mitophagy using immunofluorescence to assess the colocalization of LC3 with mitochondria in cells expressing Bnip3 and observed punctate colocalization of LC3 fluorescence with mitochondria, suggesting enhanced mitophagy. However, this observation was absent with the addition of misoprostol to Bnip3 expressing cells (Fig. 3 P, Q).

Next, we determined if Bnip3-induced mitochondrial dysfunction led to a depletion of cellular ATP stores and bioenergetic collapse. Using the FRET-based ATeam biosensor to detect cytosolic ATP levels ^45^, we observed that ATP content was significantly reduced in Bnip3-expressing H9c2 cells and that this effect was completely prevented in misoprostol treated cells (Fig. 3 R). Complementary to what we observed with acute hypoxia exposure, Bnip3-induced mitochondrial dysfunction and bioenergetic collapse in H9c2 cells translated into a significant increase in the number dead cells per field, which was also prevented with misoprostol treatment (Fig. 3 S). Together this data indicates that misoprostol is capable of inhibiting Bnip3 function and preventing mitochondrial perturbations associated with necrosis.

### 3.4. Misoprostol modulates a novel PKA phosphorylation site on Bnip3 at Thr-181

Next, we sought to determine if misoprostol was acting directly on the mitochondria or if a plasma membrane mediator was involved in this response. To do this we used isolated mitochondria from the whole rat heart, in combination with mitochondrial calcium retention capacity (CRC) and mitochondrial swelling assays that were treated directly with misoprostol or vehicle. Shown in Figure 4A and -B, misoprostol treatment had no effect on the calcium retention capacity, or the optical absorbance of isolated mitochondria treated with exogenous calcium.

**Figure 4.**
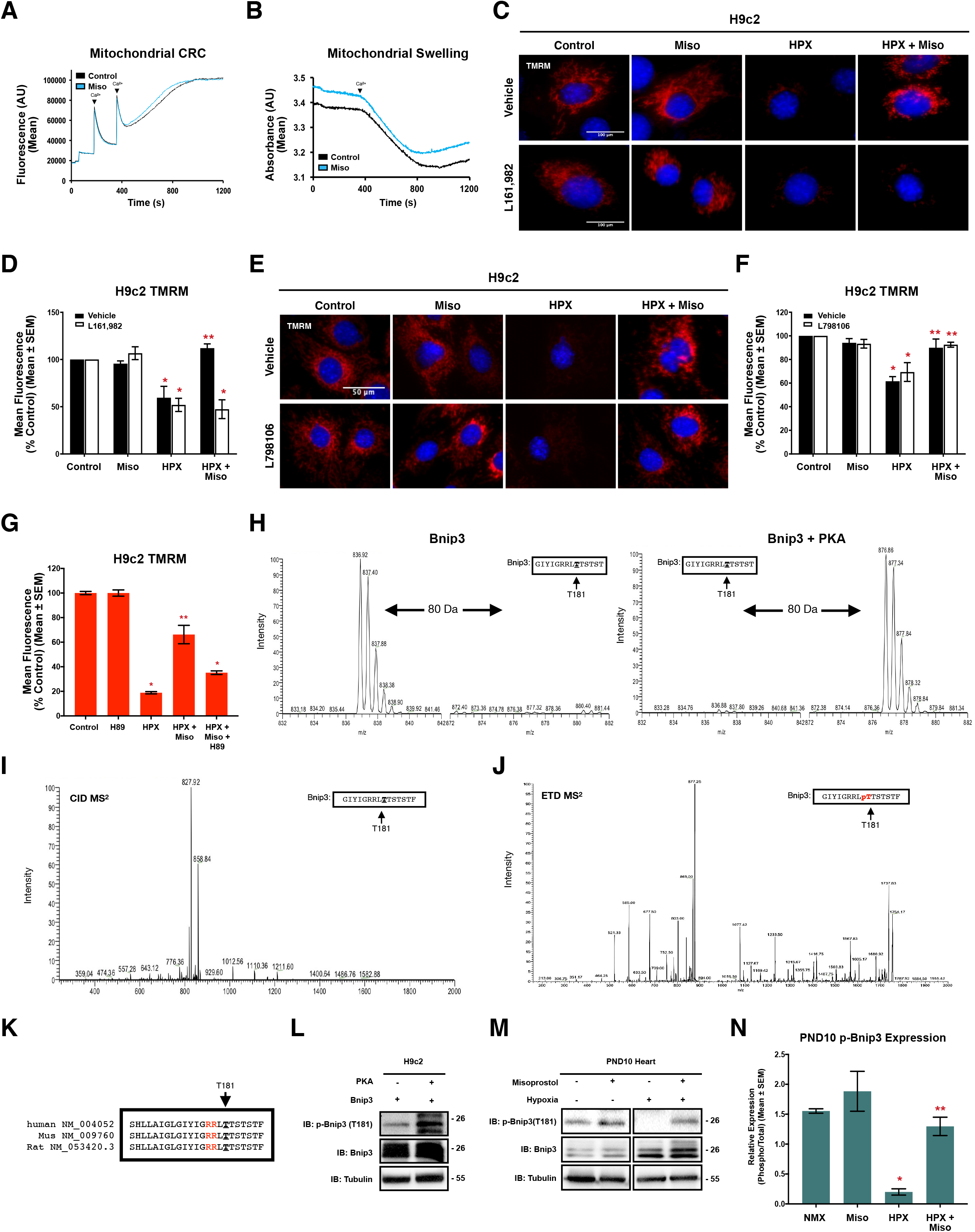
Misoprostol modulates a novel PKA phosphorylation site on Bnip3 at Thr-181. **(A)** Mitochondrial calcium retention capacity (CRC) in isolated mitochondria exposed to 10 μM misoprostol (Miso) or PBS control. **(B)** Mitochondrial swelling assay in isolated mitochondria treated as in (A). **(C)** H9c2 cells exposed to 1% O_2_ (HPX) or 21% O_2_ (control) and treated with 10 μM misoprostol (Miso) or PBS control for 24 hours. 10 μM L161,982 was also included in half of the conditions. Cells were stained with TMRM (red) and hoechst (blue) and imaged by standard fluorescence microscopy. **(D)** Quantification of cells in (C), where red fluorescent signal was normalized to cell area and quantified in 30 random fields, across 3 independent experiments. **(E)** H9c2 cells treated, stained and imaged as in (C). 1 μM L798106 was also included in half of the conditions. **(F)** Quantification of cells in (E), as in (D). **(G)** Quantification of H9c2 cells treated as in (C) with the addition of 10 μM H-89 for 24 hours. Cells were imaged as in (C) and quantified as in (D). **(H)** SIM scan of the wild-type peptide spanning the PKA site of Bnip3. The unphosphorylated peptide has a 837 m/z (z=2+) (Left), putative phosphorylation showing an increased m/z of 20 that corresponds to PO3 (M = 80.00 Da) (Right). **(I)** MS^2^ spectra following collision induced dissociation (CID) of the mass shifted ion from (h) yielding a product-ion consistent with a neutral loss of phosphate. **(J)** MS^2^ spectra following electron transfer dissociation (ETD) of a triply charged mass-shifted ion following kinase reaction (not shown). Analysis of this fragmentation spectra confirmed that threonine-181 is the preferred phosphorylation residue. **(K)** Alignment of evolutionarily conserved T181 phosphorylation site (bold and underlined), and 14-3-3 binding motifs (Red) for Bnip3 in human, mouse and rat. **(L)** Immunoblot of H9c2 cells transfected with Myc-Bnip3 and/or PKA for 16 hours. **(M)** Representative immunoblot of heart protein extracts from post-natal day (PND10) mice exposed to hypoxia (10% O_2_) ± 10 μg/kg misoprostol from PND3-10. Extracts were immunoblotted as indicated. **(N)** Phospho-Bnip3 densitometry for extracts in (M), representing an n of 3 male PND10 mouse hearts. All data are represented as mean ± S.E.M. *\*P*<0.05 compared with control, while *\*\*P*<0.05 compared with hypoxia treatment, determined by 1-way ANOVA or 2-way ANOVA where appropriate.

As misoprostol did not affect mitochondrial swelling or calcium accumulation directly, we investigated the role of prostaglandin cell surface receptors. We differentially inhibited the prostaglandin EP_3_ and EP_4_ receptors, which are both known to be enriched in the heart. Using TMRM to monitor Δ m, we applied L161,982, an EP_4_ receptor antagonist, in combination with hypoxia and misoprostol treatments. Consistent with our observations in PVNCs, misoprostol treatment prevented hypoxia-induced mitochondrial depolarization, but importantly this rescue was completely lost when the binding to the EP_4_ receptor was inhibited (Fig. 4 C, D). Conversely, EP_3_ receptor antagonism with L798106 had no effect on misoprostol’s ability to restore Δ m during hypoxic stress (Fig. 4 E, F). In addition, we expressed a plasmid-based fluorescent protein kinase A (PKA) biosensor in H9c2 cells and treated with misoprostol. The misoprostol time course shown in Supplemental Figure 5A, demonstrates that PKA activation peaks 30 minutes following misoprostol treatment, and returns to control levels within 2 hours, indicative of the type of rapid response driven by cell surface receptor activation. These results indicate that EP_4_ receptor-dependent activation of PKA might be a mechanism by which misoprostol prevents mitochondrial permeability transition. To further explore the role of PKA in misoprostol-induced protection, we used H89, a PKA inhibitor, in combination with hypoxia and misoprostol drug treatments. Consistently, hypoxia exposure significantly reduced H9c2 Δm, which was rescued with the addition of misoprostol (Fig. 4 G). However, when misoprostol treatment was combined with H89, this rescue effect was lost, indicating a role for PKA in the misoprostol-induced protection (Fig. 4 G).

To investigate whether PKA can inhibit Bnip3 function by direct phosphorylation, we performed *in silico* analysis of the mouse Bnip3 amino acid sequence, which identified two conserved potential PKA phosphorylation motifs, the first at serine (Ser)-107 and the second at threonine (Thr)-181. We engineered peptides spanning each of these regions and exposed them to *in vitro* kinase reaction with purified PKA. Following the kinase reaction, peptides were analyzed by mass spectrometry. For the peptides spanning Ser-107, no discernible peaks corresponding to the phosphorylated form of the peptide were observed (Supplement 4 A). However, for the peptides spanning Thr-181, a single ion monitoring (SIM) scan of the control peptide displayed a predominant peak at m/z of 836.92 (z=2^+^); however, following incubation with PKA the peptide showed an increased m/z of 40, representing the addition of a phosphate to the peptide (Mass = 80.00 Da) (Figure 4 H). We also evaluated if this peptide could be phosphorylated at more than one residue, but we did not detect an increased m/z of 80 (ie. 160 Da) (Supplement 4 B). Next, we analyzed the MS^2^ spectra produced by collision-induced dissociation (CID) of the mass-shifted ion with m/z = 876.86 (z=2^+^). CID typically fragments phospho-peptides resulting in the neutral loss of H_3_PO_4_, and the generation of a product-ion with a mass less 98 Da (m/z = 49 for z=2^+^). CID of the phospho-peptide spanning Thr-181 yielded a product-ion with m/z = 827.92 (delta = 48.94), indicating phosphorylation (Figure 4 I). Although these mass shifts are consistent with phosphorylation, they do not identify which of the serines or threonines are phosphorylated within the peptide. Thus, we subjected the triply charged phospho-peptide (m/z=585.27; not shown) spanning Thr-181 to electron transfer dissociation (ETD). This technique breaks peptide bonds but retains side-chain phosphorylation’s to determine specific phospho-residues. Using the MS^2^ spectra produced by ETD, Mascot software definitively identified threonine-181 of Bnip3 as the phosphorylation residue (Fig. 4 J).

To confirm that PKA phosphorylates Bnip3 in cells and *in vivo*, we used a custom phospho-specific antibody targeted to Thr-181. We co-expressed the catalytic subunit of PKA and Bnip3 in H9c2 cells and observed a marked increase in phosphorylation (p-Bnip3) (Fig. 4 L). We next exposed H9c2 cells and PVNCs to misoprostol and observed an increase in endogenous Bnip3 phosphorylation (p-Bnip3) with the Thr-181 targeted antibody (Supplement 5 B, C). Finally, to determine if Bnip3 phosphorylation is regulated during hypoxia-induced cardiac pathologies, we performed western blots on cardiac extracts from neonatal mice exposed to hypoxia for 7 days and observed a significant reduction of Bnip3 phosphorylation at Thr-181 in the hypoxic neonatal heart (Fig. 4 M, N). In addition, when the neonatal mice were both exposed to hypoxia and treated with misoprostol, Bnip3 phosphorylation was returned to control levels (Fig. 4 M, N). In addition, we evaluated Bnip3 phosphorylation in adult rodent hearts and observed a significant decrease in Bnip3 phosphorylation in the viable border zone following 4-weeks of coronary ligation (C.L) in adult Sprague Dawley rats (Supplement 3 A-C). However, during the recovery phase (8 weeks post C.L.), where the heart is overcoming the initial insult, we observed a restoration in Bnip3 phosphorylation when compared to the sham control (Supplement 3 A-C). Taken together these results imply that Bnip3 phosphorylation at Thr-181 is a regulated event during hypoxic injury *in vivo*.

### 3.5. Misoprostol inhibits Bnip3 through Thr-181

To understand the cellular role of Bnip3 phosphorylation, we first engineered a peptide spanning Thr-181 and generated a Bnip3 expression plasmid, replacing Thr-181 with an alanine residue (T181A). Using ion-trap mass spectroscopy combined with an *in vitro* kinase assay, we demonstrated that the Bnip3 mutant peptide can no longer be phosphorylated (Fig. 5 A), unlike its wild-type peptide shown previously (Fig. 4 H). To demonstrate specificity, we also engineered a peptide that replaces the threonine at position-182 with a neutral alanine and observed phosphorylation similar to the wild-type peptide (Supplement 4 C). We next employed gain-of-function transfection studies with the Bnip3-T181A construct in combination with mito-Emerald to visualize mitochondrial morphology. Similar to what was observed with the WT Bnip3 construct, expression of the T181A mutant resulted in a robust shift in mitochondrial morphology towards a fragmented and punctate phenotype (Fig. 5 B, C). However, unlike WT Bnip3, the T181A mutant was not inhibited by misoprostol treatment, and the fragmented mitochondrial morphology was retained (Fig. 5 B, C). In addition, misoprostol was not able to overcome the significant reductions in Δm that resulted from T181A expression in H9c2 cells (Fig. 5 D, E). To determine the specificity of Thr-181 as a down-stream target of misoprostol treatment, we reconstituted WT or T181A Bnip3 expression in Bnip3^-/-^ MEFs. We observed that both the WT and T181A constructs reduced mitochondrial membrane potential; however, misoprostol treatment restored Δm to control in the WT Bnip3 transfected cells but failed to significantly improve Δ m in the presence of T181A (Fig. 5 F, G).

**Figure 5.**
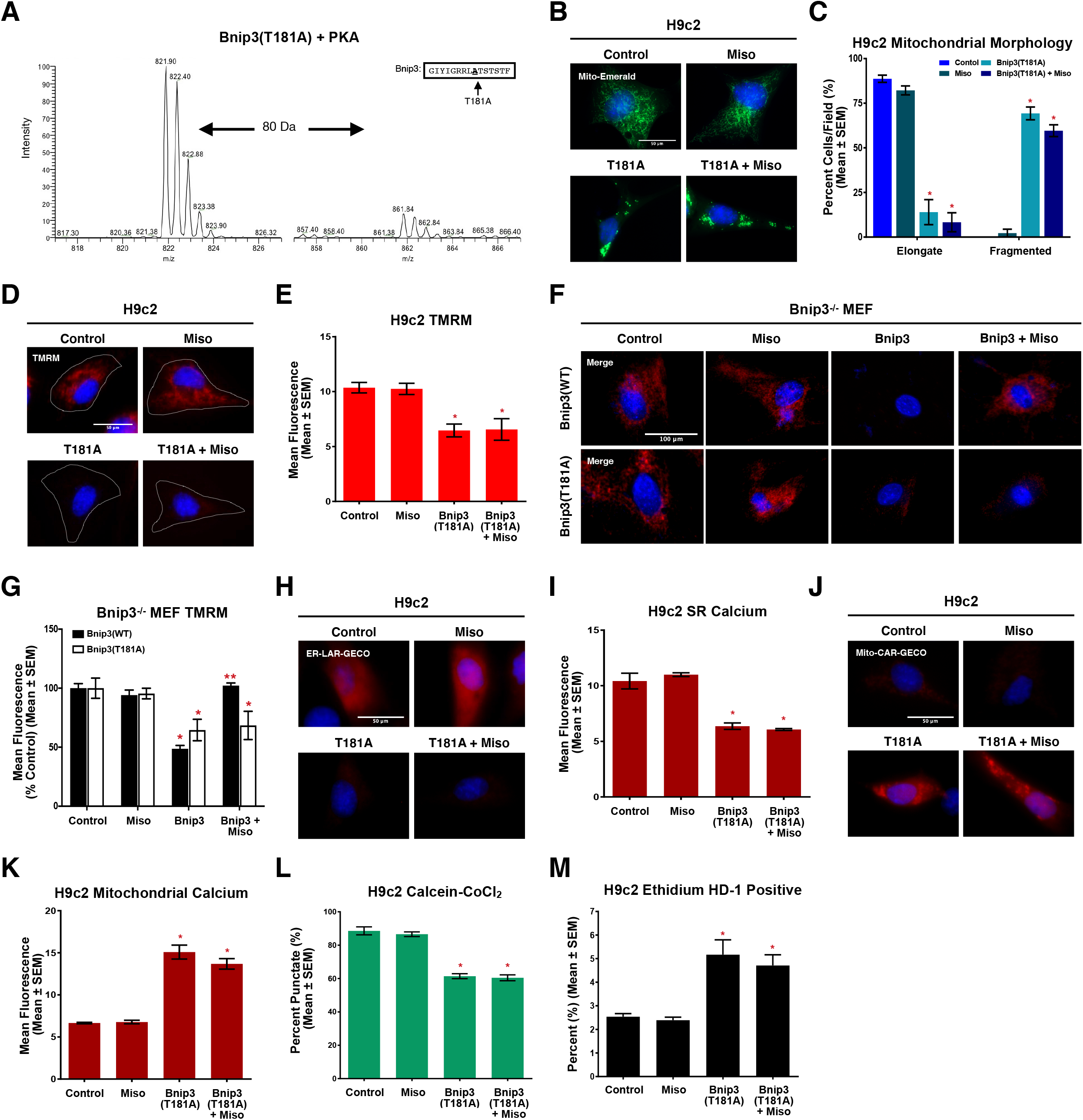
Misoprostol Inhibits Bnip3 through Thr-181. **(A)** SIM scan of a mutated peptide where the PKA site at Threonine-181 is replaced with Alanine (left). On the right, phosphorylation of this mutate peptide is negligible at the predicted m/z that corresponds to the addition of a PO_3_ (M = 80.00 Da). **(B)** H9c2 cells transfected with pcDNA3 (control) or Myc-T181A and treated with 10 μM misoprostol (Miso) or PBS control for 16 hours. Mito-Emerald (green) was included in all conditions to show transfected cells and mitochondrial morphology. Cells were stained with hoechst (blue) and imaged by standard fluorescence microscopy. **(C)** Quantification of cells in (B), where the number of cells with elongated and fragmented mitochondria are expressed as a percentage of all transfected cells in 30 random fields, across 3 independent experiments. **(D)** H9c2 cells treated as in (B). CMV-GFP (outline) was included in all conditions to indicate transfected cells Cells were stained with TMRM (red) and hoechst (blue) and imaged by standard fluorescence microscopy. **(E)** Quantification of cells in (D), where red fluorescent signal was normalized to cell area and quantified in 30 random fields, across 3 independent experiments. **(F)** Bnip3^-/-^ mouse embryonic fibroblasts (MEFs) treated as in (B) where either Myc-Bnip3(WT) or Myc-Bnip3(T181A) was transfected in. Cells were stained with TMRM (red) and hoechst (blue) and imaged by standard fluorescence microscopy. **(G)** Quantification of cells in (F), where red fluorescent signal was normalized to cell area and quantified in 15 random fields, across 3 independent experiments. **(H)** H9c2 cells treated as in (B). ER-LAR-GECO (red) was included in all conditions to indicate ER calcium content. Cells were stained with hoechst (blue) and imaged by standard fluorescence microscopy. **(I)** Quantification of cells in (H) as in (E) in 30 random fields, across 3 independent experiments. **(J)** H9c2 cells treated as in (B). Mito-CAR-GECO (red) was included in all conditions to indicate mitochondrial calcium content. Cells were stained with hoechst (blue) and imaged by standard fluorescence microscopy. **(K)** Quantification of cells in (J) as in (E) in 30 random fields, across 3 independent experiments. **(L)** Quantification of H9c2 cells treated as in (B) and CMV-ds.RED was included in all conditions to indicate transfected cells. Cells were stained with hoechst (blue) and calcein-AM quenched by cobalt chloride (CoCl_2_, 5 μM) to assess permeability transition. Quantification was done by calculating the percentage of cells with mitochondrial puncta in 30 random fields, across 3 independent experiments. **(M)** Quantification of H9c2 cells treated as in (B). Live cells were stained with calcein-AM (green), and necrotic cells were stained with ethidium homodimer-1 (red) and are expressed as percent (%) dead in 30 random fields, across 3 independent experiments. All data are represented as mean ± S.E.M. *\*P*<0.05 compared with control, while *\*\*P*<0.05 compared with hypoxia treatment, determined by 1-way ANOVA or 2-way ANOVA where appropriate.

When we investigated that underlying calcium phenomena, we observed that like WT, T181A expression shifted calcium away from the ER and into the mitochondria. However, unlike WT, misoprostol was unable to prevent this calcium movement in the presence of the T181A mutant (Fig. 5 H-K). Furthermore, using calcein-CoCl_2_ and Live/Dead imaging, we observed that the T181A mutant was sufficient to drive mitochondrial permeability transition and cell death, which could not be prevented with misoprostol drug treatment (Fig. 5 L, M). These data demonstrate that phosphorylation of Bnip3 at Thr-181 is necessary for misoprostol to inhibit Bnip3 function in multiple cell models.

### 3.6. Thr-181 Phosphorylation Retains Bnip3 in the cytosol

To determine how phosphorylation at Thr-181 inhibits Bnip3 function, we expressed a mitochondrial matrix-targeted green fluorescent plasmid in H9c2 cells, and performed immunofluorescence for Bnip3 following exposure to hypoxia and/or misoprostol. As shown in Figure 6 A and B, confocal microscopy revealed that at baseline there is very little interaction between the organelle-targeted fluorophores and Bnip3; however, when H9c2 cells were exposed to hypoxia the colocalization coefficient is increased by 116.3% at the mitochondria Interestingly, we observed that this organellar localization was abrogated with the addition of concurrent misoprostol treatment (Fig. 6 A, B). Using a similar approach, we assessed the colocalization between Bnip3 and mitochondrial Opa1 protein in the neonatal heart. Like in H9c2 cells, at baseline there was very little interaction between the two proteins, which was significantly increased in response to hypoxia exposure (Fig. 6 C). However, this observed hypoxia-induced colocalization between Bnip3 and Opa1 was disrupted in the presence of misoprostol in the neonatal heart (Fig. 6 C). Additionally, when we used an SR/ER targeted green fluorescent plasmid in H9c2 cells to assess potential interactions between Bnip3 and the ER, we observed that exposure to hypoxia increased the colocalization of the two targets by 381.9% (Fig. 6 D). Interestingly, we observed that this organellar localization was also abrogated with the addition of misoprostol treatment (Fig. 6 D). Next, we determined the subcellular localization of phosphorylated Bnip3 through subcellular fractionation studies. We observed that Bnip3 is predominantly localized to the mitochondria, and to a lesser extent at the ER, while phosphorylated Bnip3 is retained in the cytosol (Fig. 6 E). These findings are consistent with our observation in Figure 4M, showing that Bnip3 phosphorylation is decreased during hypoxia, and restored by misoprostol treatment, which also inhibits Bnip3-dependent mitochondrial perturbations.

**Figure 6.**
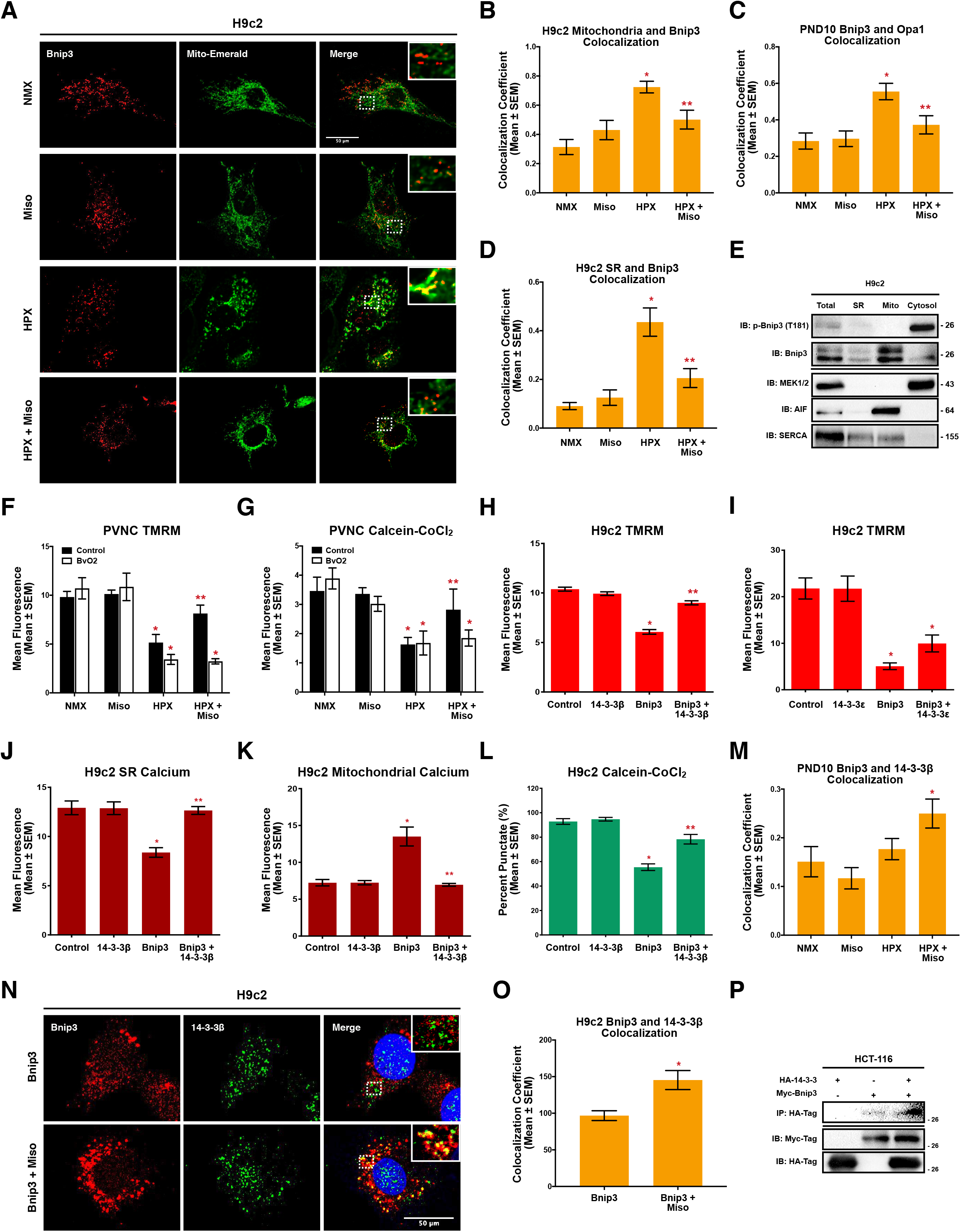
Thr-181 Phosphorylation Retains Bnip3 in the cytosol. **(A)** H9c2 cells treated with 10 μ misoprostol (Miso) ± 1% O_2_ (HPX) for 24 hours. Myc-Bnip3 and Mito-Emerald (green) were included in each condition to visualize localization. Cells were fixed, stained with hoechst (blue), and immunofluorescence was performed using a Myc-tag primary antibody (Red). Cells were then imaged by standard confocal microscopy. **(B)** Quantification of cells in (A), where colocalization coefficient was calculated for 30 cells per condition across 10 random fields. **(C)** Quantification of immunofluorescence in PND10 hearts exposed to hypoxia (10% O_2_) ± 10 μg/kg misoprostol from PND3-10. Hearts were probed for Bnip3 (Red), Opa1 (Green), and stained with DAPI (Blue). Hearts were imaged by standard confocal microscopy and the colocalization coefficient was calculated in 20 fields per condition (n=4 animals/conditions). **(D)** Quantification of H9c2 cells treated as in (A). ER-Emerald (green) was transfected in all conditions. Cells were fixed, stained with hoechst (blue), and immunofluorescence was performed using a Myc-tag primary antibody (Red). Cells were then imaged by standard confocal microscopy. Colocalization coefficient was calculated for 30 cells per condition across 10 random fields. **(E)** Fractionation of control treated H9c2. Protein extracts were fractionated and immunoblotted, as indicated. **(F)** Quantification of primary ventricular neonatal cardiomyocytes (PVNCs) treated with 10 μM misoprostol (Miso) ± 1% O_2_ (HPX) for 24 hours. 5 μM BvO2 was included in half of the conditions to inhibit 14-3-3 protein activity. Cells were stained with TMRM (red) and hoechst (blue) and imaged by standard fluorescence microscopy. Red fluorescent signal was normalized to cell area and quantified in 20 random fields, across 2 independent experiments. **(G)** Quantification of PVNC’s treated as in (E). Cells were stained with hoechst (blue) and calcein-AM quenched by cobalt chloride (CoCl_2_, 5 μM) to assess permeability transition, and imaged by standard fluorescence microscopy. The percentage of cells with mitochondrial puncta was calculated in 20 random fields, across 2 independent experiments. **(H)** Quantification of H9c2 cells transfected with pcDNA3 (control) or Myc-Bnip3 with and without HA-14-3-3β. CMV-GFP (outline) was included in all conditions to indicate transfected cells Cells were stained with TMRM (red) and hoechst (blue) and imaged by standard fluorescence microscopy. Red fluorescent signal was normalized to cell area and quantified in 30 random fields, across 3 independent experiments. **(I)** Quantifcation of H9c2s treated as in (G), with and without 14-3-3ε. Cells were stained and imaged as in (H) and quantified as an (I) across 30 random fields, in 3 independent experiments. **(J)** Quantification of H9c2’s treated as in (H). ER-LAR-GECO (red) was included in all conditions to indicate ER calcium content. Cells were stained and imaged as in (H). Red fluorescent signal was normalized to cell are in 30 random fields, across 3 independent experiments. **(K)** Quantification of H9c2’s treated as in (H). Mito-CAR-GECO (red) was included in all conditions to indicate mitochondrial calcium content. Quantification performed as in (J) in 30 random fields, across 3 independent experiments. **(L)** Quantification of H9c2’s treated as in (H), and CMV-ds.RED was included in all conditions to indicate transfected cells. Cells were stained with hoechst (blue) and calcein-AM quenched by cobalt chloride (CoCl_2_, 5 μM) to assess permeability transition. Where the percentage of cells with mitochondrial puncta was calculated in 30 random fields, across 3 independent experiments. **(M)** Quantification of immunofluorescence in PND10 hearts exposed to hypoxia (10% O_2_) ± 10 μg/kg misoprostol from PND3-10. Hearts were probed for Bnip3 (Red), 14-3-3β (Green), and stained with DAPI (Blue). Hearts were imaged by standard confocal microscopy and the colocalization coefficient was calculated in 20 fields per condition (n=4 animals/conditions). **(N)** H9c2 cells transfected with Myc-Bnip3 ± 10 μM misoprostol (Miso) 18 hours. Cells were fixed, stained with hoechst (blue), and immunofluorescence was performed using a Myc-tag (Red), and 14-3-3β (Green). Cells were then imaged by standard confocal microscopy. **(O)** Quantification of cells in (N), where colocalization coefficient was calculated for 30 cells per condition across 10 random fields. **(P)** Co-immunoprecipitation of HCT-116 cells expressing HA-14-3-3 and Myc-Bnip3. Proteins were pulled down with Myc and probed for HA-tag. Immunoblot was probed as indicated. All data are represented as mean ± S.E.M. *\*P*<0.05 compared with control, while *\*\*P*<0.05 compared with hypoxia treatment, determined by 1-way ANOVA or 2-way ANOVA where appropriate.

*In silico* analysis of Bnip3 predicted that Thr-181 lies within a conserved interacting domain of the molecular chaperone family 14-3-3. While 14-3-3 family members are able to identify a number of sequences, Bnip3 possess the classical RxxpTx motif (Bnip3: RRLpTT), which are commonly found within PKA and CaMKII phosphorylation sites (See Fig. 4K for alignment). As certain 14-3-3 family members are known interactors with Bcl-2 proteins ^55–59^, and we recently determined that PKA-dependent phosphorylation of Nix (Bnip3L) increases its interaction with 14-3-3β ^60^, we investigated the role of these molecular chaperones as a mechanism by which misoprostol inhibits Bnip3 function. Using hypoxia and misoprostol exposed PVNCs, we applied BvO2, a pan-14-3-3 inhibitor, and assessed mitochondrial membrane potential using TMRM. We observed that misoprostol’s ability to rescue of Δ m was prevented with 14-3-3 inhibition (Fig. 6 F). We observed similar results when we used calcein-CoCl_2_ to visualize MPT, where misoprostol treatment prevented hypoxia-induced permeability transition, in the absence but not the presence of BvO2 (Fig. 6 G).

We were further interested in determining which 14-3-3 family member is involved in this mechanism. Based on previous data from our group, which demonstrated that 14-3-3β traffics phosphorylated Nix from the mitochondria and ER/SR in skeletal muscle cell lines, we began by investigating this 14-3-3 family member in the cardiomyocyte ^60^. Using gain of function transfection studies where we expressed Bnip3 and 14-3-3β, alone and in combination, in H9c2 cells. TMRM staining revealed that like misoprostol, 14-3-3β was able to rescue Bnip3-induced mitochondrial depolarizations (Fig. 6 H). Similar experiments were conducted using 14-3-3ε, which was unable to restore mitochondrial membrane potential, demonstrating some degree of isoform specificity (Fig. 6 I). Next, we determined that 14-3-3β expression is sufficient to prevent the ER calcium depletion and mitochondrial calcium accumulation that is triggered by Bnip3 expression (Fig. 6 J, K). Using the calcein-CoCl_2_ method, we also evaluated how these calcium events were affecting MPT. Similar to our previous results, Bnip3 significantly increases the number of H9c2 cells experiencing MPT in each field, reducing the number of cells with distinct mitochondrial puncta. However, consistent with the mitochondrial calcium results, when we combined Bnip3 and 14-3-3β expression, MPT was prevented, and cells returned to their normal punctate phenotype (Fig. 6 L).

Given this data indicating a potential functional interaction between 14-3-3β and Bnip3, we were next interested in determining if there was a physical interaction between the two proteins. We started by performing immunofluorescence targeting both Bnip3 and 14-3-3β following exposure to hypoxia and/or misoprostol in the neonatal heart. This approach, in combination with confocal microscopy, revealed that at baseline there was very little colocalization between the two proteins; likely due to the relatively low expression of Bnip3 in the normoxic conditions. However, the combination of hypoxia exposure and misoprostol drug treatments significantly increased the colocalization coefficient between Bnip3 and 14-3-3β (Fig. 6 M). To validate this finding and overcome the challenges associated with altered Bnip3 expression levels between normoxic and hypoxic conditions, we overexpressed both Bnip3 and 14-3-3β in H9c2 cells and performed confocal immunofluorescence. As shown in Figure 6 N and O, the addition of misoprostol increased the degree of colocalization by more than 45%. Next, we co-expressed HA-14-3-3β and myc-Bnip3 in HCT-116 cells and performed a co-immunoprecipitation with the HA antibody. Using this approach, we observed a marked increase in detectable myc-Bnip3 when the two proteins were expressed together, indicative of a physical interaction between the two proteins (Fig. 6 P). Collectively, this data indicates that misoprostol promotes Bnip3 trafficking away from the mitochondria and ER, resulting in p-Bnip3 almost exclusively residing in the cytosol. In addition, Bnip3 strongly colocalizes with mitochondrial Opa1 during hypoxic conditions, but this is attenuated by misoprostol treatment in favour of the inhibitory function of the 14-3-3β chaperone.

### 3.7 Bnip3 ablation prevents hypoxia-induced contractile dysfunction and necroinflammation in the neonatal heart

To determine if a direct link existed between hypoxia-induced alterations in contractile function and Bnip3 protein expression *in vivo*, we returned to our mouse model of neonatal hypoxia this time using mice harboring a genetic deletion of Bnip3 ^37^. Using this approach in combination with transthoracic echocardiography, we observed that hypoxia induced significant contractile dysfunction in wild-type neonatal animals, including reductions in ejection fraction (EF), and alterations in left ventricular filling between heart beats (E’/A’) (Fig. 7 A-C). When we assessed Bnip3-null mice under that same conditions, there were no alterations in contractile function at baseline (normoxia) and the loss of Bnip3 conferred protection against hypoxia-induced derangements in contraction, preventing reductions in both EF and E’/A’ (Fig. 7 A-C). These results phenocopy what we observed using misoprostol drug treatments during neonatal hypoxia. At the same time, when we investigated the subcellular distribution of HMGB1 in these PND10 hearts, we found that when WT animals were exposed to hypoxia this resulted a marked decrease in HMGB1 nuclear localization (Fig, 7 D). However, when we investigated the distribution of HMGB1 in the hearts of hypoxia-exposed Bnip3^-/-^ mice, we observed that the inflammatory protein was retained in the nucleus (Fig. 7 D).

**Figure 7.**
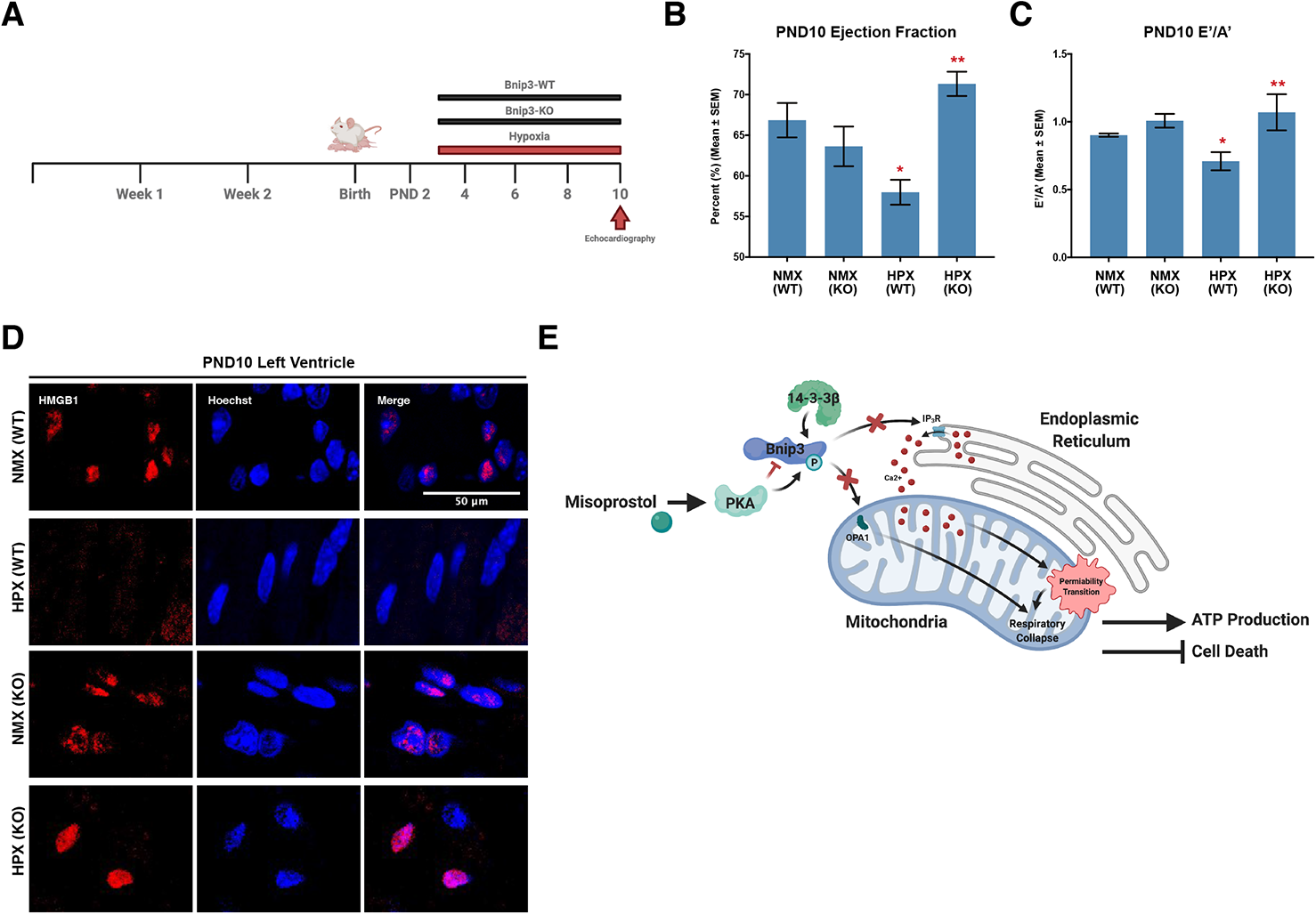
Bnip3 ablation prevents hypoxia-induced contractile dysfunction and necroinflammation in the neonatal heart. **(A)** Post-natal day 2 (PND2) Bnip3-WT and Bnip3-null mice are exposed to hypoxia (10% O_2_) from PND3-10, hearts were imaged and collected on PND10. **(B)** Ejection fraction and **(C)** E’/A’ ratio for PND10 animals treated as in (A), in 3-5 mice per group, as determined by transthoracic echocardiography. **D)** PND10 hearts treated as in (A) and stained with DAPI (Blue) and probed for high mobility group box 1 (HMGB1, red). Hearts were imaged via confocal microscopy. **(E)** Proposed mechanism by which misoprostol inhibits Bnip3 at the mitochondria and ER to prevent necrotic cell death and necroinflammation. All data are represented as mean ± S.E.M. *\*P*<0.05 compared with WT control, while *\*\*P*<0.05 compared with WT hypoxia treatment, determined by 1-way ANOVA.

## 4. Discussion

According to the World Health Organization, systemic hypoxia is the leading complication associated with perinatal emergencies, contributing to the majority of neonatal deaths within the first week of life. Sustained hypoxia is also a known driver of functional alterations in the neonatal heart, resulting in contractile impairment, resistance to inotropic therapy, and end-organ perfusion defects ^61–67^. While until now the mechanisms have remained unclear, in this study we provide *in vivo* and cell-based evidence, including data from human iPSC-derived cardiomyocytes, that hypoxia-induced mitochondrial permeability transition, bioenergetic collapse, and necroinflammation contributes to neonatal cardiac dysfunction. Through several lines of investigation, we established that this pathology is contingent on Bnip3 expression. We further demonstrate that Bnip3 is sufficient to alter mitochondrial calcium homeostasis and oxidative-phosphorylation failure, ultimately resulting in mitochondrial permeability transition and necrotic cell death, and is necessary for the *in vivo* release of the pro-inflammatory alarmin/DAMP HMGB1 in the neonatal heart. We also demonstrate that Bnip3 function can be pharmacologically modulated by misoprostol leading to activation of EP_4_ receptors, PKA signalling, and direct phosphorylation at threonine-181, which localizes Bnip3 to the cytosol enhancing the colocalization with 14-3-3β and preventing the colocalization with Opa1 (See Fig. 7 E for overview).

The results presented in this report serve to unify previous reports demonstrating the deleterious role of Bnip3 and its pro-death C-terminal transmembrane (TM) domain. Through detailed evaluations of the role of Bnip3 at ER/SR, we demonstrate changes in subcellular calcium localization. Previous data suggests that the TM domain of Bnip3 directly interacts with Bcl-2, which is traditionally associated with inhibiting the IP_3_R, resulting in calcium release from the ER ^14^. This Bnip3-induced ER calcium is quickly buffered by the mitochondria through VDAC and MCU directly in the mitochondrial matrix ^18–20^. These previous studies further demonstrate that mitochondrial calcium drives a loss of membrane potential and respiration, ROS production, MPT and ultimately a caspase-independent necrosis ^14, 17, 68^. However, Bnip3 is known as a dual-regulator of cell death, where it also inserts through the outer mitochondrial membrane and uses its TM domain to interact with the dynamin related protein, Opa1 ^16, 69, 70^. While Opa1 is traditionally associated with maintaining cristae structure, efficient organization of ETC complexes, and mitochondrial fusion, disruption and/or genetic deletion of Opa1 results in ETC dysfunction, mitochondrial fragmentation and cell death ^15, 71, 72^. Additionally, work by Rikka *et al.* demonstrated that cardiomyocyte-specific overexpression of Bnip3 enhances mitochondrial protease activity, resulting in the degradation of complex-1 (NADH ubiquinone oxidoreductase) and −4 (cytochrome c oxidase), suppressing respiratory activity, while enhancing mitochondrial fragmentation ^29^. ETC dysfunction is further tied to the overproduction of mitochondrial reactive oxygen species, which propagates an influx of calcium into the matrix and synergizes with MPT ^73, 74^. Together these studies demonstrate that by altering mitochondrial function and calcium homeostasis in two very different ways, Bnip3 displays overlapping and redundant functions to depress energy production and promote necrotic cell death in the heart.

Consistent with these studies, we show that in the context of neonatal pathological hypoxic signaling, Bnip3 expression in the heart drives mitochondrial dysfunction and cell death. By directly measuring SR/ER calcium content this work advances our understanding of what Bnip3 is doing at the organellar level, and supports the notion that Bnip3 triggers SR/ER calcium release that is buffered by the mitochondria. We further build on the findings of these past studies that link mitochondrial calcium accumulation with ROS production alongside the induction of MPT, bioenergetic collapse and necrosis both in cultured cardiomyocytes and *in vivo*. Importantly the work presented here also demonstrates that these pathways can be pharmacologically modulated through misoprostol-induced EP4 activation, which work from us and others, has been shown to be cardioprotective ^10, 34, 53^. While a previous study has demonstrated that Bnip3 phosphorylation inhibits its interactions with Opa1, we provide mechanistic evidence both *in vivo* and in cardiomyocytes that misoprostol activates the EP4 receptor and PKA, resulting in an inhibitory phosphorylation of Bnip3’s TM domain at Thr-181. Furthermore, based on the known roles of the 14-3-3 family of molecular chaperones, we propose a mechanism by which 14-3-3β translocates Bnip3 to the cytosol, which likely prevents interactions with factors at ER and mitochondrial, including Bcl-2 and Opa1, respectively ^55–59^.

Taken together the results presented in this study elevate the role of Bnip3 in the neonatal heart, and strongly implicate it as a critical regulator of mitochondrial calcium homeostasis that when upregulated by hypoxia drives calcium into the mitochondria and a necroinflammatory phenotype. Additionally, the data in this preclinical study builds on the accumulating evidence that misoprostol directly regulates Bnip3 function, with potential meaningful implications for neonatal and adult hypoxia-induced cardiac pathologies, and stem cell-based cardiac therapies where promoting cardiomyocyte survival would be of benefit.

## Funding and Acknowledgments

This work was supported by the Natural Science and Engineering Research Council (NSERC) Canada, through Discovery Grants to J.W.G. and A.R.W.. This work is supported by Heart and Stroke Foundation of Canada (HSFC) Grants to J.W.G. and G.M.H. V.W.D and C.A.D. are supported by CIHR. G.M.H is a Canada Research Chair in Cardiolipin Metabolism. M.D.M. and N.S. are supported by studentships from the Children’s Hospital Foundation of Manitoba and Research Manitoba, M.D.M. received support from the DEVOTION research cluster. A.S. is supported by grants from SSHRC New Frontiers in Research Fund, Research Manitoba, Children’s Hospital Research Institute of Manitoba, and Children’s Hospital Foundation. We thank Dr. ‘s Bill Diehl-Jones and Yan Hai for their support during the preliminary phase of this study. We also thank Farhana Begum from the Histomorphology and Ultrastructural Imaging Core in the Department of Human Anatomy and Cell Science at the University of Manitoba for her technical expertise that made this work possible. This work was conducted at the Children’s Hospital Research Institute of Manitoba.

## Conflict of Interest

None.

## Author Contributions

M.D.M. and J.W.G. conceptualized the study. M.D.M. and J.W.G. were responsible for writing the paper. M.D.M. designed and conducted most of the investigations and was responsible for data curation. M.D.M. was also responsible for formal analysis and data visualization. N.S. conducted both the Seahorse XF-24 investigation and cellular/cardiac ATP assays, including data curation and formal analysis. D.C. conducted the Bnip3 qRT-PCR investigations. B.X. conducted PND10 *in vivo* transthoracic echocardiography. C.R. and J.W.G designed and conducted ion-trap mass spec studies. J.M.K. and A.M. conducted isolated mitochondrial studies. I.M.D. and S.R. designed and conducted M.I. model. J.W.G., and A.R.W. were responsible for funding acquisition. All authors reviewed the results, edited, and approved the final version of the manuscript.

**Supplement 3. Phosphorylation of Bnip3 at Thr-181 is altered in adult hypoxia-induced pathologies. (A)** Representative immunoblot of heart protein extracts from Sprague Dawley rats subjected to left coronary artery ligation, or sham operation as a control. Following 4-weeks of recovery, the viable infarct border-zone was harvested from the left ventricle. Extracts were immunoblotted for phospho-Bnip3 expression. **(B)** Densitometry analysis for extracts in (A), representing an N of 4 animals per condition. **(C)** Representative immunoblot of heart protein extracts from Sprague Dawley rats treated as in (A) and viable infarct border-zone was harvested 8-weeks of recovery. Extracts were immunoblotted for phospho-Bnip3 expression. All data are represented as mean ± S.E.M. *\*P*<0.05 compared with control, while *\*\*P*<0.05 compared with hypoxia treatment, determined by 1-way ANOVA or 2-way ANOVA where appropriate.

**Supplement 4. Characterization of Bnip3 phosphorylation by mass spectroscopy. (A)** SIM scan of the peptide spanning the predicted S107 PKA site of Bnip3. The unphosphorylated peptide has a 625 m/z (z=2+), which is not changed in the presence of PKA. **(B)** SIM scan of the wild-type peptide spanning the T181 PKA site of Bnip3. The unphosphorylated peptide has a 837 m/z (z=2+) which is shifted by phosphorylation showing an increased m/z of 20 that corresponds to a single PO3 (M = 80.00 Da) and not a double phosphorylation (M=160.00 Da). **(C)** SIM scan of a mutated peptide where the PKA site at Threonine-182 is replaced with Alanine. The unphosphorylated peptide has a 837 m/z (z=2+), while putative phosphorylation shows an increased m/z of 20 that corresponds to PO3 (M = 80.00 Da), demonstrating no effect of T182A on PKA’s ability to phosphorylate Bnip3.

**Supplement 5. Regulation of Bnip3 phosphorylation by misoprostol and PKA. (A)** Quantification of H9c2 cells transfected with pPHT-PKA and treated with 10 μM misoprostol for 0.5-h, 1-h, 2-h and 4-h. Cells were imaged by standard fluorescence microscopy and the ratio of green (active) to red (inactive) fluorescent signal was measured, normalized to cell area, and quantified in 10 random fields. **(B)** Immunoblot for phospho-Bnip3(T181) in protein extracts from H9c2 cells treated with 10 μM misoprostol or PBS control for 16 hours. **(C)** Immunoblot for phospho-Bnip3(T181) in protein extracts from PVNC’s treated with 10 μM misoprostol or PBS control for 30 minutes. All data are represented as mean ± S.E.M. *\*P*<0.05 compared with control, while *\*\*P*<0.05 compared with hypoxia treatment, determined by 1-way ANOVA or 2-way ANOVA where appropriate.

**Supplement 6. Misoprostol alters Bnip3’s interactions with Opa1 and 14-3-3β. (A)** PND10 hearts from mice exposed to hypoxia (10% O_2_) ± 10 μg/kg misoprostol from PND3-10, stained with DAPI (Blue) and probed for Bnip3 (Red), and Opa1 (green). Hearts were imaged via confocal microscopy. **(B)** PND10 hearts from mice treated as in (A) and stained with DAPI (Blue) and probed for Bnip3 (Red), and 14-3-3β (green). Hearts were imaged via confocal microscopy.

## Notes

### Competing Interest Statement

The authors have declared no competing interest.

### Summary of Updates

This version of the manuscript includes new in vivo and cell culture data regarding the regulation of Bnip3 and Opa1 through Misoprostol treatment and the molecular chaperone 14-3-3 beta.

## References

1. UNICEF, WHO, World Bank. Levels & Trends in Child Mortality: Report 2020 : Estimates Developed by the UN Inter-Agency Group for Child Mortality Estimation. WHO Geneva: United Nations Children’s Fund; 2020.

2. Luu TM, Katz SL, Leeson P, Thebaud B, Nuyt A-M. Preterm birth: risk factor for early-onset chronic diseases. Can Med Assoc J 2015;188:736–746.

3. Vannucci SJ. Hypoxia-ischemia in the immature brain. J Exp Biol 2004;207:3149–3154.

4. Shastri AT, Samarasekara S, Muniraman H, Clarke P. Cardiac troponin i concentrations in neonates with hypoxic-ischaemic encephalopathy. Acta Paediatr Int J Paediatr 2012;101:26–29.

5. Armstrong K, Franklin O, Sweetman D, Molloy EJ. Cardiovascular dysfunction in infants with neonatal encephalopathy. Arch Dis Child 2012;97:372–375.

6. Greer SN, Metcalf JL, Wang Y, Ohh M. The updated biology of hypoxia-inducible factor. EMBO J 2012;31:2448–2460.

7. Chaudhuri RD, Banerjee D, Banik A, Sarkar S. Severity and duration of hypoxic stress differentially regulates HIF-1α-mediated cardiomyocyte apoptotic signaling milieu during myocardial infarction. Arch Biochem Biophys 2020;690:108430.

8. Carmeliet P, Dor Y, Herbert J-M, Fukumura D, Brusselmans K, Dewerchin M, Neeman M, Bono F, Abramovitch R, Maxwell P, Koch CJ, Ratcliffe P, Moons L, Jain RK, Collen D, Keshet E. Role of HIF-1α in hypoxia-mediated apoptosis, cell proliferation and tumour angiogenesis. Nature 1998;394:485–490.

9. Gustafsson ÅB. Bnip3 as a dual regulator of mitochondrial turnover and cell death in the myocardium. Pediatr Cardiol 2011;32:267–274.

10. Field JT, Martens MD, Mughal W, Hai Y, Chapman D, Hatch GM, Ivanco TL, Diehl-Jones W, Gordon JW. Misoprostol regulates Bnip3 repression and alternative splicing to control cellular calcium homeostasis during hypoxic stress. Cell Death Discov 2018;4:98–98.

11. Kubli DA, Ycaza JE, Gustafsson ÅB. Bnip3 mediates mitochondrial dysfunction and cell death through Bax and Bak. Biochem J 2007;405:407–415.

12. Azad MB, Chen Y, Henson ES, Cizeau J, McMillan-Ward E, Israels SJ, Gibson SB. Hypoxia induces autophagic cell death in apoptosis-competent cells through a mechanism involving BNIP3. Autophagy 2008;4:195–204.

13. Gordon JW, Shaw JA, Kirshenbaum LA. Multiple facets of NF-kB in the heart: To be or not to NF-kB. Circ Res 2011;108:1122–1132.

14. Ray R, Chen G, Vande Velde C, Cizeau J, Park JH, Reed JC, Gietz RD, Greenberg AH. BNIP3 Heterodimerizes with Bcl-2/Bcl-XL and Induces Cell Death Independent of a Bcl-2 Homology 3 (BH3) Domain at Both Mitochondrial and Nonmitochondrial Sites. J Biol Chem 2000;275:1439–1448.

15. Liu KE, Frazier WA. Phosphorylation of the BNIP3 C-terminus inhibits mitochondrial damage and cell death without blocking autophagy. PLoS ONE 2015;10:1–28.

16. Landes T, Emorine LJ, Courilleau D, Rojo M, Belenguer P, Arnauné-Pelloquin L. The BH3-only Bnip3 binds to the dynamin Opa1 to promote mitochondrial fragmentation and apoptosis by distinct mechanisms. EMBO Rep 2010;11:459–465.

17. Zhang L, Li L, Liu H, Borowitz JL, Isom GE. BNIP3 mediates cell death by different pathways following localization to endoplasmic reticulum and mitochondrion. FASEB J 2009;23:3405–3414.

18. Rapizzi E, Pinton P, Szabadkai G, Wieckowski MR, Vandecasteele G, Baird G, Tuft RA, Fogarty KE, Rizzuto R. Recombinant expression of the voltage-dependent anion channel enhances the transfer of Ca2+ microdomains to mitochondria. J Cell Biol 2002;159:613–624.

19. Baughman JM, Perocchi F, Girgis HS, Plovanich M, Belcher-Timme CA, Sancak Y, Bao XR, Strittmatter L, Goldberger O, Bogorad RL, Koteliansky V, Mootha VK. Integrative genomics identifies MCU as an essential component of the mitochondrial calcium uniporter. Nature 2011;476:341–345.

20. Chaudhuri D, Sancak Y, Mootha VK, Clapham DE. MCU encodes the pore conducting mitochondrial calcium currents. eLife 2013;2:e00704.

21. Karch J, Kwong JQ, Burr AR, Sargent MA, Elrod JW, Peixoto PM, Martinez-Caballero S, Osinska H, Cheng EH-Y, Robbins J, Kinnally KW, Molkentin JD. Bax and Bak function as the outer membrane component of the mitochondrial permeability pore in regulating necrotic cell death in mice. eLife 2013;2:e00772.

22. Giorgio V, Stockum S von, Antoniel M, Fabbro A, Fogolari F, Forte M, Glick GD, Petronilli V, Zoratti M, Szabo I, Lippe G, Bernardi P. Dimers of mitochondrial ATP synthase form the permeability transition pore. Proc Natl Acad Sci 2013;110:5887–5892.

23. Izzo V, Bravo-San Pedro JM, Sica V, Kroemer G, Galluzzi L. Mitochondrial Permeability Transition: New Findings and Persisting Uncertainties. Trends Cell Biol 2016;26:655–667.

24. Mughal W, Martens M, Field J, Chapman D, Huang J, Rattan S, Hai Y, Cheung KG, Kereliuk S, West AR, Cole LK, Hatch GM, Diehl-Jones W, Keijzer R, Dolinsky VW, Dixon IM, Parmacek MS, Gordon JW. Myocardin regulates mitochondrial calcium homeostasis and prevents permeability transition. Cell Death Differ 2018;25:1732–1748.

25. Whelan RS, Kaplinskiy V, Kitsis RN. Cell Death in the Pathogenesis of Heart Disease: Mechanisms and Significance. Annu Rev Physiol 2010;72:19–44.

26. Baines CP, Kaiser RA, Purcell NH, Blair NS, Osinska H, Hambleton MA, Brunskill EW, Sayen MR, Gottlieb RA, Dorn GW, Robbins J, Molkentin JD. Loss of cyclophilin D reveals a critical role for mitochondrial permeability transition in cell death. Nature 2005;434:658–662.

27. Nakagawa T, Shimizu S, Watanabe T, Yamaguchi O, Otsu K, Yamagata H, Inohara H, Kubo T, Tsujimoto Y. Cyclophilin D-dependent mitochondrial permeability transition regulates some necrotic but not apoptotic cell death. Nature 2005;434:652–658.

28. Kwong JQ, Davis J, Baines CP, Sargent MA, Karch J, Wang X, Huang T, Molkentin JD. Genetic deletion of the mitochondrial phosphate carrier desensitizes the mitochondrial permeability transition pore and causes cardiomyopathy. Cell Death Differ 2014;21:1209–1217.

29. Rikka S, Quinsay MN, Thomas RL, Kubli DA, Zhang X, Murphy AN, Gustafsson ÅB. Bnip3 impairs mitochondrial bioenergetics and stimulates mitochondrial turnover. Cell Death Differ 2011;18:721–731.

30. Baker PE, Fahey JV, Munck A. Prostaglandin inhibition of T-cell proliferation is mediated at two levels. Cell Immunol 1981;61:52–61.

31. Lanefelt F, Ullberg M, Jondal M, Fredholm BB. PGE1 and prostacyclin suppression of NK-cell mediated cytotoxicity and its relation to cyclic AMP. Med Biol 1983;61:324–330.

32. Syeda MM, Jing X, Mirza RH, Yu H, Sellers RS, Chi Y. Prostaglandin Transporter Modulates Wound Healing in Diabetes by Regulating Prostaglandin-Induced Angiogenesis. Am J Pathol 2012;181:334–346.

33. Martens MD, Fernando AS, Gordon JW. A New Trick for an Old Dog? Myocardial-Specific Roles for Prostaglandins as Mediators of Ischemic Injury and Repair. Am J Physiol-Heart Circ Physiol 2021;ajpheart.00872.2020.

34. Bryson TD, Gu X, Khalil RM, Khan S, Zhu L, Xu J, Peterson E, Yang X-P, Harding P. Overexpression of prostaglandin E2 EP4 receptor improves cardiac function after myocardial infarction. J Mol Cell Cardiol 2018;118:1–12.

35. Bryson TD, Pandrangi TS, Khan SZ, Xu J, Pavlov TS, Ortiz PA, Peterson EL, Harding P. The Deleterious Role of the Prostaglandin E2 EP3 Receptor in Angiotensin II Hypertension. Am J Physiol-Heart Circ Physiol 2020;318:H867–H882.

36. Kubasiak LA, Hernandez OM, Bishopric NH, Webster KA. Hypoxia and acidosis activate cardiac myocyte death through the Bcl-2 family protein BNIP3. Proc Natl Acad Sci 2002;99:12825–12830.

37. Diwan A, Krenz M, Syed FM, Wansapura J, Ren X, Koesters AG, Li H, Kirshenbaum LA, Hahn HS, Robbins J, Jones WK, Dorn GW. Inhibition of ischemic cardiomyocyte apoptosis through targeted ablation of Bnip3 restrains postinfarction remodeling in mice. J Clin Invest 2007;117:2825–2833.

38. Ward NL, Moore E, Noon K, Spassil N, Keenan E, Ivanco TL, LaManna JC. Cerebral angiogenic factors, angiogenesis, and physiological response to chronic hypoxia differ among four commonly used mouse strains. J Appl Physiol Bethesda Md 1985 2007;102:1927–1935.

39. Wallace MG, Hartle KD, Snow WM, Ward NL, Ivanco TL. Effect of hypoxia on the morphology of mouse striatal neurons. Neuroscience 2007;147:90–96.

40. Dixon IM, Lee SL, Dhalla NS. Nitrendipine binding in congestive heart failure due to myocardial infarction. Circ Res 1990;66:782–788.

41. Ghavami S, Cunnington RH, Gupta S, Yeganeh B, Filomeno KL, Freed DH, Chen S, Klonisch T, Halayko AJ, Ambrose E, Singal R, Dixon IMC. Autophagy is a regulator of TGF-β1-induced fibrogenesis in primary human atrial myofibroblasts. Cell Death Dis 2015;6:e1696.

42. Dolinsky VW, Morton JS, Oka T, Robillard-Frayne I, Bagdan M, Lopaschuk GD, Des Rosiers C, Walsh K, Davidge ST, Dyck JRB. Calorie Restriction Prevents Hypertension and Cardiac Hypertrophy in the Spontaneously Hypertensive Rat. Hypertension 2010;56:412–421.

43. Planchon TA, Gao L, Milkie DE, Davidson MW, Galbraith JA, Galbraith CG, Betzig E. Rapid three-dimensional isotropic imaging of living cells using Bessel beam plane illumination. Nat Methods 2011;8:417–423.

44. Wu J, Liu L, Matsuda T, Zhao Y, Rebane A, Drobizhev M, Chang YF, Araki S, Arai Y, March K, Hughes TE, Sagou K, Miyata T, Nagai T, Li WH, Campbell RE. Improved orange and red Ca2+ indicators and photophysical considerations for optogenetic applications. ACS Chem Neurosci 2013;4:963–972.

45. Kotera I, Iwasaki T, Imamura H, Noji H, Nagai T. Reversible Dimerization of *Aequorea victoria* Fluorescent Proteins Increases the Dynamic Range of FRET-Based Indicators. ACS Chem Biol 2010;5:215–222.

46. Ding Y, Li J, Enterina JR, Shen Y, Zhang I, Tewson PH, Mo GCH, Zhang J, Quinn AM, Hughes TE, Maysinger D, Alford SC, Zhang Y, Campbell RE. Ratiometric biosensors based on dimerization-dependent fluorescent protein exchange. Nat Methods 2015;12:195–198.

47. Mishra P, Carelli V, Manfredi G, Chan DC. Proteolytic cleavage of Opa1 stimulates mitochondrial inner membrane fusion and couples fusion to oxidative phosphorylation. Cell Metab 2014;19:630–641.

48. Diehl-Jones W, Archibald A, Gordon JW, Mughal W, Hossain Z, Friel JK. Human Milk Fortification Increases Bnip3 Expression Associated With Intestinal Cell Death In Vitro. J Pediatr Gastroenterol Nutr 2015;61:583–590.

49. Moghadam AR, Silva Rosa SC da, Samiei E, Alizadeh J, Field J, Kawalec P, Thliveris J, Akbari M, Ghavami S, Gordon JW. Autophagy modulates temozolomide-induced cell death in alveolar Rhabdomyosarcoma cells. Cell Death Discov 2018;4:52.

50. Seshadri N, Jonasson ME, Hunt KL, Xiang B, Cooper S, Wheeler MB, Dolinsky VW, Doucette CA. Uncoupling protein 2 regulates daily rhythms of insulin secretion capacity in MIN6 cells and isolated islets from male mice. Mol Metab 2017;6:760–769.

51. Karch J, Bround MJ, Khalil H, Sargent MA, Latchman N, Terada N, Peixoto PM, Molkentin JD. Inhibition of mitochondrial permeability transition by deletion of the ANT family and CypD. Sci Adv 2019;5:eaaw4597.

52. Mughal W, Nguyen L, Pustylnik S, Silva Rosa SC da, Piotrowski S, Chapman D, Du M, Alli NS, Grigull J, Halayko AJ, Aliani M, Topham MK, Epand RM, Hatch GM, Pereira TJ, Kereliuk S, McDermott JC, Rampitsch C, Dolinsky VW, Gordon JW. A conserved MADS-box phosphorylation motif regulates differentiation and mitochondrial function in skeletal, cardiac, and smooth muscle cells. Cell Death Dis 2015;6:e1944–e1944.

53. Martens MD, Field JT, Seshadri N, Day C, Chapman D, Keijzer R, Doucette CA, Hatch GM, West AR, Ivanco TL, Gordon JW. Misoprostol attenuates neonatal cardiomyocyte proliferation through Bnip3, perinuclear calcium signaling, and inhibition of glycolysis. J Mol Cell Cardiol 2020;146:19–31.

54. Galluzzi L, Vitale I, Aaronson SA, Abrams JM, Adam D, Agostinis P, Alnemri ES, Altucci L, Amelio I, Andrews DW, Annicchiarico-Petruzzelli M, Antonov AV, Arama E, Baehrecke EH, Barlev NA, Bazan NG, Bernassola F, Bertrand MJM, Bianchi K, Blagosklonny MV, Blomgren K, Borner C, Boya P, Brenner C, Campanella M, Candi E, Carmona-Gutierrez D, Cecconi F, Chan FK- M, Chandel NS, et al. Molecular mechanisms of cell death: recommendations of the Nomenclature Committee on Cell Death 2018. Cell Death Differ 2018;25:486–541.

55. Datta SR, Katsov a, Hu L, Petros a, Fesik SW, Yaffe MB, Greenberg ME. 14-3-3 proteins and survival kinases cooperate to inactivate BAD by BH3 domain phosphorylation. Mol Cell 2000;6:41–51.

56. Masters SC, Fu H. 14-3-3 Proteins Mediate an Essential Anti-apoptotic Signal. J Biol Chem 2001;276:45193–45200.

57. Petosa C, Masters SC, Bankston LA, Pohl J, Wang B, Fu H, Liddington RC. 14-3-3z Binds a Phosphorylated Raf Peptide and an Unphosphorylated Peptide Via Its Conserved Amphiphatic Groove. J Biol Chem 1998;273:16305–16310.

58. Tzivion G, Avruch J. 14-3-3 Proteins: Active Cofactors in Cellular Regulation by Serine/Threonine Phosphorylation. J Biol Chem 2002;277:3061–3064.

59. Tan Y, Demeter MR, Ruan H, Comb MJ. BAD Ser-155 phosphorylation regulates BAD/Bcl-XL interaction and cell survival. J Biol Chem 2000;275:25865–25869.

60. Silva Rosa SC da, Martens MD, Field JT, Nguyen L, Kereliuk SM, Hai Y, Chapman D, Diehl-Jones W, Aliani M, West AR, Thliveris J, Ghavami S, Rampitsch C, Dolinsky VW, Gordon JW. BNIP3L/Nix-induced mitochondrial fission, mitophagy, and impaired myocyte glucose uptake are abrogated by PRKA/PKA phosphorylation. Autophagy 2020;1–16.

61. Miall-Allen VM, Vries LS de, Whitelaw AG. Mean arterial blood pressure and neonatal cerebral lesions. Arch Dis Child 1987;62:1068–1069.

62. Goldstein RF, Thompson RJ, Oehler JM, Brazy JE. Influence of acidosis, hypoxemia, and hypotension on neurodevelopmental outcome in very low birth weight infants. Pediatrics 1995;95:238–243.

63. Baker CFW, Barks JDE, Engmann C, Vazquez DM, Neal CR, Schumacher RE, Bhatt-Mehta V. Hydrocortisone administration for the treatment of refractory hypotension in critically ill newborns. J Perinatol 2008;28:412–419.

64. Joynt C, Cheung P-Y. Treating Hypotension in Preterm Neonates With Vasoactive Medications. Front Pediatr 2018;6:86.

65. Joynt C, Cheung P-Y. Cardiovascular Supportive Therapies for Neonates With Asphyxia - A Literature Review of Pre-clinical and Clinical Studies. Front Pediatr 2018;6:363.

66. Carr H, Cnattingius S, Granath F, Ludvigsson JF, Edstedt Bonamy AK. Preterm Birth and Risk of Heart Failure Up to Early Adulthood. J Am Coll Cardiol 2017;69:2634–2642.

67. Cox DJ, Edwards AD, Hajnal JV, Durighel G, Price AN, Broadhouse KM, Groves AM, Finnemore AE. Cardiovascular magnetic resonance of cardiac function and myocardial mass in preterm infants: a preliminary study of the impact of patent ductus arteriosus. J Cardiovasc Magn Reson 2014;16:1–9.

68. Vande Velde C, Cizeau J, Dubik D, Alimonti J, Brown T, Israels S, Hakem R, Greenberg AH. BNIP3 and genetic control of necrosis-like cell death through the mitochondrial permeability transition pore. Mol Cell Biol 2000;20:5454–5468.

69. Chen Y, Lewis W, Diwan A, Cheng EH-Y, Matkovich SJ, Dorn GW. Dual autonomous mitochondrial cell death pathways are activated by Nix/BNip3L and induce cardiomyopathy. Proc Natl Acad Sci 2010;107:9035–9042.

70. Pereira RO, Tadinada SM, Zasadny FM, Oliveira KJ, Pires KMP, Olvera A, Jeffers J, Souvenir R, Mcglauflin R, Seei A, Funari T, Sesaki H, Potthoff MJ, Adams CM, Anderson EJ, Abel ED. OPA1 deficiency promotes secretion of FGF21 from muscle that prevents obesity and insulin resistance. EMBO J 2017;36:2126–2145.

71. Frezza C, Cipolat S, Martins de Brito O, Micaroni M, Beznoussenko GV, Rudka T, Bartoli D, Polishuck RS, Danial NN, De Strooper B, Scorrano L. OPA1 Controls Apoptotic Cristae Remodeling Independently from Mitochondrial Fusion. Cell 2006;126:177–189.

72. Cogliati S, Frezza C, Soriano ME, Varanita T, Quintana-Cabrera R, Corrado M, Cipolat S, Costa V, Casarin A, Gomes LC, Perales-Clemente E, Salviati L, Fernandez-Silva P, Enriquez JA, Scorrano L. Mitochondrial Cristae Shape Determines Respiratory Chain Supercomplexes Assembly and Respiratory Efficiency. Cell 2013;155:160–171.

73. Lopez-Fabuel I, Le Douce J, Logan A, James AM, Bonvento G, Murphy MP, Almeida A, Bolaños JP. Complex I assembly into supercomplexes determines differential mitochondrial ROS production in neurons and astrocytes. Proc Natl Acad Sci 2016;113:13063–13068.

74. Koopman WJH, Verkaart S, Visch HJ, Emst-de Vries S van, Nijtmans LGJ, Smeitink JAM, Willems PHGM. Human NADH:ubiquinone oxidoreductase deficiency: radical changes in mitochondrial morphology? Am J Physiol-Cell Physiol 2007;293:C22–C29.

