## Supplementary material for "Misoprostol Treatment Prevents Hypoxia-Induced Cardiac Dysfunction Through a 14-3-3 and PKA regulatory motif on Bnip3": Table 1

**Supplement 1. Complete list of relative Ct values for mitochondrial energy metabolism qRT-PCR array in PND10 heart samples (n=3 animals/condition), identified by mitochondrial complex.**

| Complex | Target | Array ID | NMX | HPX | HPX + MISO |
|---|---|---|---|---|---|
| Complex 1 | Ndufa10 | Mm00600325_m1 | 1.000000 | 0.197397 | 0.225888 |
| | Ndufs4 | Mm00656176_m1 | 1.000000 | 0.251290 | 0.626054 |
| | Ndufa6 | Mm01303455_g1 | 1.000000 | 0.446720 | 0.560977 |
| | Ndufab1 | Mm01137654_g1 | 1.000000 | 0.456153 | 0.530460 |
| | Ndufa11 | Mm01236867_g1 | 1.000000 | 0.462330 | 0.876922 |
| | Ndufb3 | Mm00835179_g1 | 1.000000 | 0.477876 | 5.086023 |
| | Lhpp | Mm01269918_m1 | 1.000000 | 0.867248 | 2.086091 |
| | Ndufb7 | Mm00788005_s1 | 1.000000 | 0.883464 | 3.577282 |
| | Ndufb9 | Mm00612543_m1 | 1.000000 | 0.963204 | 4.114191 |
| | Ndufs7 | Mm01144210_m1 | 1.000000 | 0.993365 | 1.360459 |
| | Ndufa5 | Mm01165335_m1 | 1.000000 | 1.053928 | 2.232583 |
| | Ndufa3 | Mm01329704_g1 | 1.000000 | 1.119881 | 2.969799 |
| | Ndufs3 | Mm01329746_g1 | 1.000000 | 1.270824 | 0.955209 |
| | Ndufs6 | Mm02529639_u1 | 1.000000 | 1.272835 | 1.198779 |
| | Ndufs5 | Mm02600127_g1 | 1.000000 | 1.384180 | 3.076444 |
| | Ndufv3 | Mm01345700_m1 | 1.000000 | 1.870935 | 0.721520 |
| | Ndufc2 | Mm04213119_s1 | 1.000000 | 2.068060 | 1.442927 |
| | Ndufs2 | Mm00467603_g1 | 1.000000 | 2.454698 | 0.991675 |
| | Ndufs1 | Mm00523640_m1 | 1.000000 | 3.377395 | 2.718098 |
| | Ndufb5 | Mm00452592_m1 | 1.000000 | 5.968531 | 2.784598 |
| Complex 2 | Sdhd | Mm00546511_m1 | 1.000000 | 0.652135 | 0.805184 |
| Complex 3 | Cyc1 | Mm00470540_m1 | 1.000000 | 1.051789 | 1.276995 |
| | Uqcrq | Mm00772880_m1 | 1.000000 | 1.118767 | 0.921522 |
| | Uqcr11 | Mm00824470_g1 | 1.000000 | 1.519776 | 0.988861 |
| | Uqcrc1 | Mm00445911_m1 | 1.000000 | 2.289718 | 0.946879 |
| | Uqcrh | Mm00835199_g1 | 1.000000 | 8.377721 | 0.874225 |
| Complex 4 | Cox11 | Mm01615963_g1 | 1.000000 | 0.078951 | 0.971551 |
| | Cox4i1 | Mm01250094_m1 | 1.000000 | 0.676532 | 0.852358 |
| | Cox5a | Mm00432638_m1 | 1.000000 | 0.860288 | 0.942858 |
| | Cox7b | Mm00835076_g1 | 1.000000 | 1.087345 | 0.839040 |
| | Cox7a2 | Mm00438299_m1 | 1.000000 | 1.423320 | 2.855101 |
| | Cox8c | Mm01325374_m1 | 1.000000 | 3.089582 | 0.000000 |
| ATP Synthase | Atp5g3 | Mm01334541_g1 | 1.000000 | 0.078880 | 0.130007 |
| | Atp5d | Mm00502864_m1 | 1.000000 | 0.677803 | 4.668294 |
| | Atp5a1 | Mm00431960_m1 | 1.000000 | 4.229016 | 0.646722 |
