## Supplementary material for "Misoprostol Treatment Prevents Hypoxia-Induced Cardiac Dysfunction Through a 14-3-3 and PKA regulatory motif on Bnip3": Table 2

**Supplement 2. Complete list of relative Ct values for cell death pathway finder qRT-PCR array in PND10 heart samples (n=3 animals/condition), identified by cell death pathway.**

| Pathway | Target | Array ID | NMX | HPX | HPX + MISO |
|---|---|---|---|---|---|
| Apoptosis | Mcl1 | Mm00725832_s1 | 1.00000 | 0.09085 | 0.23385 |
| | Rab25 | Mm00444175_m1 | 1.00000 | 0.26157 | 0.34519 |
| | Cflar | Mm01255578_m1 | 1.00000 | 0.39314 | 0.71528 |
| | Birc2 | Mm00431811_m1 | 1.00000 | 0.81076 | 4.67332 |
| | Bcl2 | Mm00477631_m1 | 1.00000 | 2.16225 | 0.43035 |
| | Xiap | Mm01311594_mH | 1.00000 | 5.13869 | 5.39754 |
| | Ywhaz | Mm01158417_g1 | 1.00000 | 8.17255 | 1.46103 |
| | Bcl2l11 | Mm00437796_m1 | 1.00000 | 0.33395 | 1.44004 |
| | Casp1 | Mm00438023_m1 | 1.00000 | 0.34537 | 1.73104 |
| | Apaf1 | Mm01223702_m1 | 1.00000 | 0.67758 | 1.14831 |
| | Cd40 | Mm00441891_m1 | 1.00000 | 2.10615 | 1.11862 |
| | Casp2 | Mm00432314_m1 | 1.00000 | 2.52140 | 5.98441 |
| | Fas | Mm01204974_m1 | 1.00000 | 3.69722 | 4.33228 |
| | Traf2 | Mm00801978_m1 | 1.00000 | 3.90931 | 1.41652 |
| | Dffa | Mm00438410_m1 | 1.00000 | 6.43212 | 0.40410 |
| Autophagy | Esr1 | Mm00433149_m1 | 1.00000 | 0.08536 | 0.12262 |
| | Gaa | Mm00484581_m1 | 1.00000 | 0.17815 | 0.36807 |
| | Akt1 | Mm01331626_m1 | 1.00000 | 0.21520 | 0.26158 |
| | Casp3 | Mm01195085_m1 | 1.00000 | 0.23869 | 1.15156 |
| | Pik3c3 | Mm00619489_m1 | 1.00000 | 0.34942 | 0.36887 |
| | Atg5 | Mm00504340_m1 | 1.00000 | 0.53311 | 5.37482 |
| | Irgm1 | Mm00492596_m1 | 1.00000 | 0.59373 | 0.37320 |
| | Map1lc3a | Mm00458725_g1 | 1.00000 | 0.66540 | 7.28435 |
| | Atg7 | Mm00512209_m1 | 1.00000 | 3.30256 | 0.76212 |
| | Becn1 | Mm01265461_m1 | 1.00000 | 3.62075 | 5.66199 |
| | Atp6v1g2 | Mm01159330_g1 | 1.00000 | 4.97828 | 6.60147 |
| | Snca | Mm01188700_m1 | 1.00000 | 5.44339 | 5.97623 |
| | Atg16l1 | Mm00513085_m1 | 1.00000 | 5.48796 | 1.04288 |
| | Mapk8 | Mm00489514_m1 | 1.00000 | 8.34948 | 9.06761 |
| Necrosis | Bmf | Mm00506773_m1 | 1.00000 | 0.16720 | 4.62077 |
| | Cyld | Mm00557599_m1 | 1.00000 | 0.51276 | 8.72694 |
| | Kcnip1 | Mm01189526_m1 | 1.00000 | 0.19440 | 0.67120 |
| | Dpysl4 | Mm00496436_m1 | 1.00000 | 0.23837 | 2.38527 |
| | Spata2 | Mm00468039_g1 | 1.00000 | 0.31416 | 1.10926 |
| | Ccdc103 | Mm01248720_m1 | 1.00000 | 0.42559 | 1.86050 |
| | Parp2 | Mm00456462_m1 | 1.00000 | 0.55469 | 2.04393 |
| | Parp1 | Mm01321084_m1 | 1.00000 | 0.59457 | 0.56596 |
| | Tmem57 | Mm00550603_m1 | 1.00000 | 0.65448 | 1.23867 |
| | Dennd4a | Mm00768872_m1 | 1.00000 | 5.50594 | 1.00824 |
| Inflammation | Il12b | Mm01288992_m1 | 1.00000 | 0.27653 | 0.01620 |
| | Edn1 | Mm00438656_m1 | 1.00000 | 0.45629 | 0.40834 |
| | Il1b | Mm00434228_m1 | 1.00000 | 0.49674 | 0.54534 |
| | Il15 | Mm00434210_m1 | 1.00000 | 2.05814 | 4.37305 |
| | Cd4 | Mm00442754_m1 | 1.00000 | 4.82734 | 2.44139 |
| | Il12a | Mm00434165_m1 | 1.00000 | 10.19767 | 0.00000 |
| | Il17a | Mm00439619_m1 | 1.00000 | 10.33179 | 0.00000 |
