## Supplemental figure for "Misoprostol Treatment Prevents Hypoxia-Induced Cardiac Dysfunction Through a 14-3-3 and PKA regulatory motif on Bnip3"

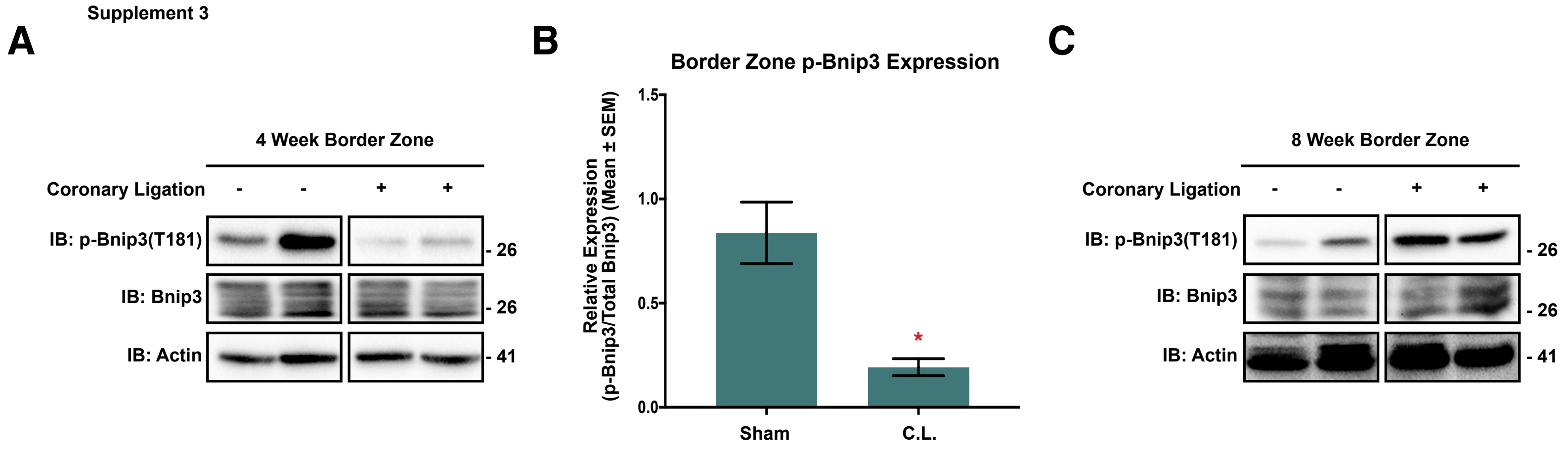


**Supplement 3. Phosphorylation of Bnip3 at Thr-181 is altered in adult hypoxia-induced pathologies. (A)** Representative immunoblot of heart protein extracts from Sprague Dawley rats subjected to left coronary artery ligation, or sham operation as a control. Following 4-weeks of recovery, the viable infarct border-zone was harvested from the left ventricle. Extracts were immunoblotted for phospho-Bnip3 expression. **(B)** Densitometry analysis for extracts in (A), representing an N of 4 animals per condition. **(C)** Representative immunoblot of heart protein extracts from Sprague Dawley rats treated as in (A) and viable infarct border-zone was harvested 8-weeks of recovery. Extracts were immunoblotted for phospho-Bnip3 expression. All data are represented as mean ± S.E.M. *$P$<0.05 compared with control determined by Students T-Test.
