## Supplemental figure for "Misoprostol Treatment Prevents Hypoxia-Induced Cardiac Dysfunction Through a 14-3-3 and PKA regulatory motif on Bnip3"

Supplement 4

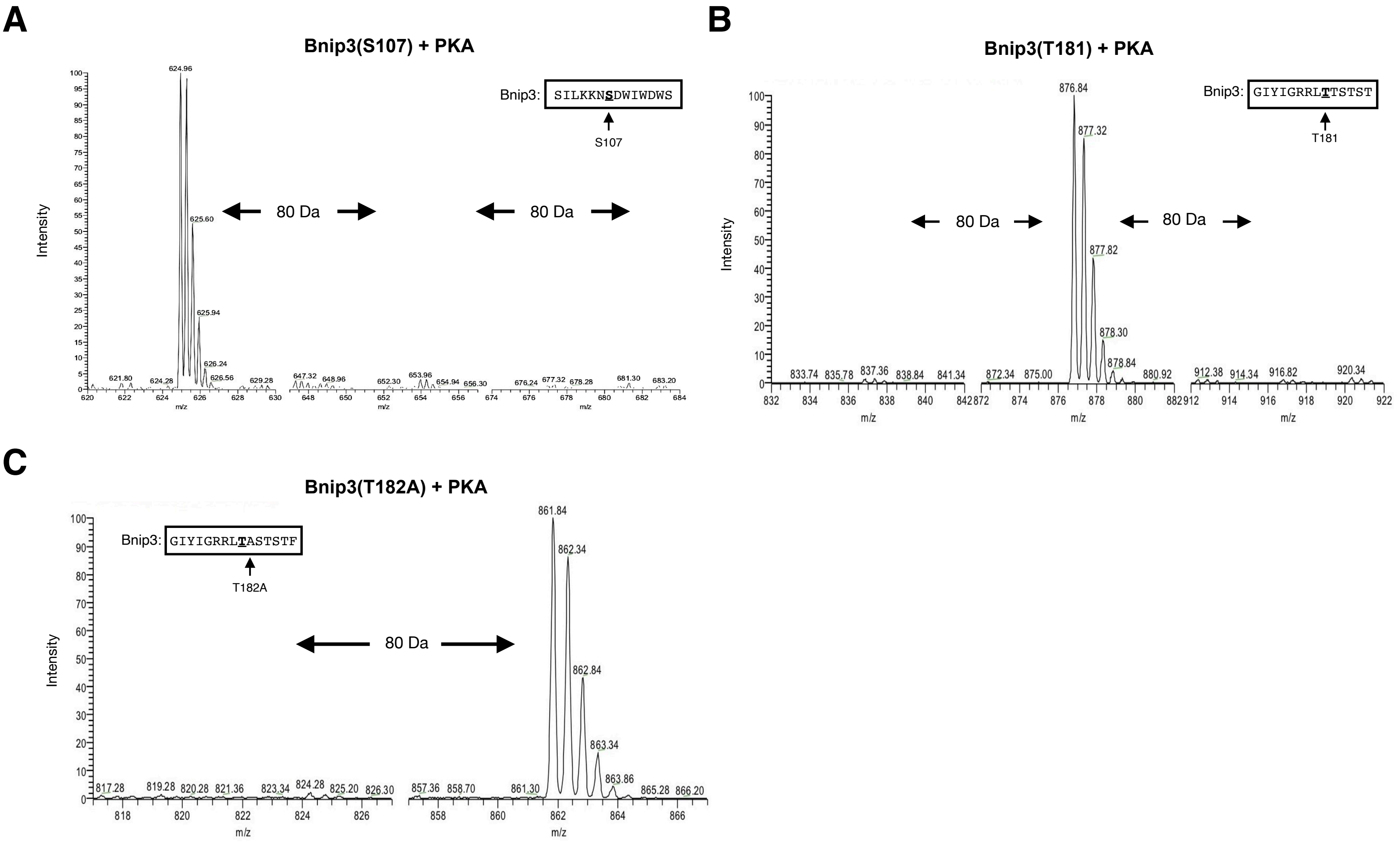


**Supplement 4. Characterization of Bnip3 phosphorylation by mass spectroscopy. (A)** SIM scan of the peptide spanning the predicted S107 PKA site of Bnip3. The unphosphorylated peptide has a 625 m/z (z=2+), which is not changed in the presence of PKA. **(B)** SIM scan of the wild-type peptide spanning the T181 PKA site of Bnip3. The unphosphorylated peptide has a 837 m/z (z=2+) which is shifted by phosphorylation showing an increased m/z of 20 that corresponds to a single PO3 (M = 80.00 Da) and not a double phosphorylation (M=160.00 Da). **(C)** SIM scan of a mutated peptide where the PKA site at Threonine-182 is replaced with Alanine. The unphosphorylated peptide has a 837 m/z (z=2+), while putative phosphorylation shows an increased m/z of 20 that corresponds to PO3 (M = 80.00 Da), demonstrating no effect of T182A on PKA's ability to phosphorylate Bnip3.
