## Supplemental figure for "Misoprostol Treatment Prevents Hypoxia-Induced Cardiac Dysfunction Through a 14-3-3 and PKA regulatory motif on Bnip3"

Supplement 5

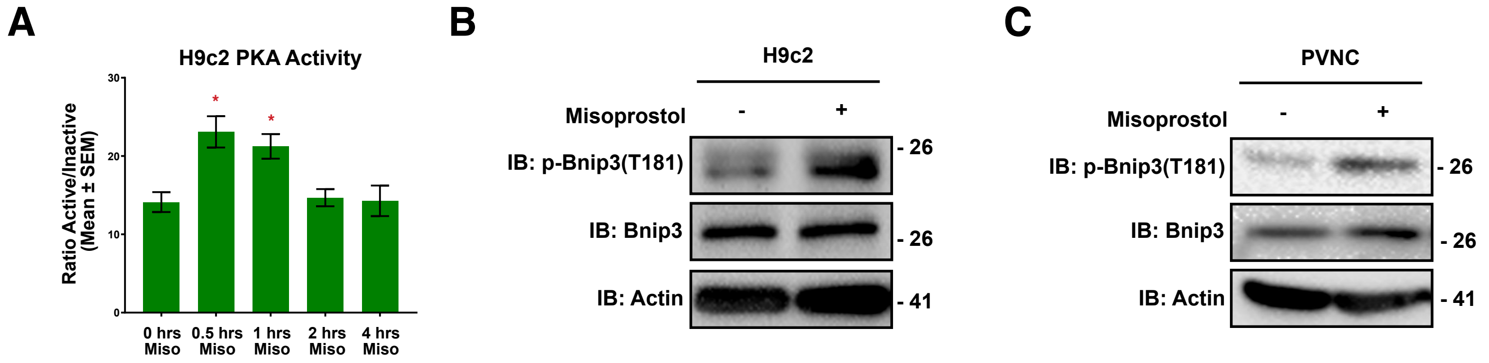


**Supplement 5. Regulation of Bnip3 phosphorylation by misoprostol and PKA. (A)** Quantification of H9c2 cells transfected with pPHT-PKA and treated with 10 μM misoprostol for 0.5-h, 1-h, 2-h and 4-h. Cells were imaged by standard fluorescence microscopy and the ratio of green (active) to red (inactive) fluorescent signal was measured, normalized to cell area, and quantified in 10 random fields. **(B)** Immunoblot for phospho-Bnip3(T181) in protein extracts from H9c2 cells treated with 10 μM misoprostol or PBS control for 16 hours. **(C)** Immunoblot for phospho-Bnip3(T181) in protein extracts from PVNC's treated with 10 μM misoprostol or PBS control for 30 minutes. All data are represented as mean ± S.E.M. **P*<0.05 compared with control, while ***P*<0.05 compared with hypoxia treatment, determined by 1-way ANOVA or 2-way ANOVA where appropriate.
