## Supplemental figure for "Misoprostol Treatment Prevents Hypoxia-Induced Cardiac Dysfunction Through a 14-3-3 and PKA regulatory motif on Bnip3"

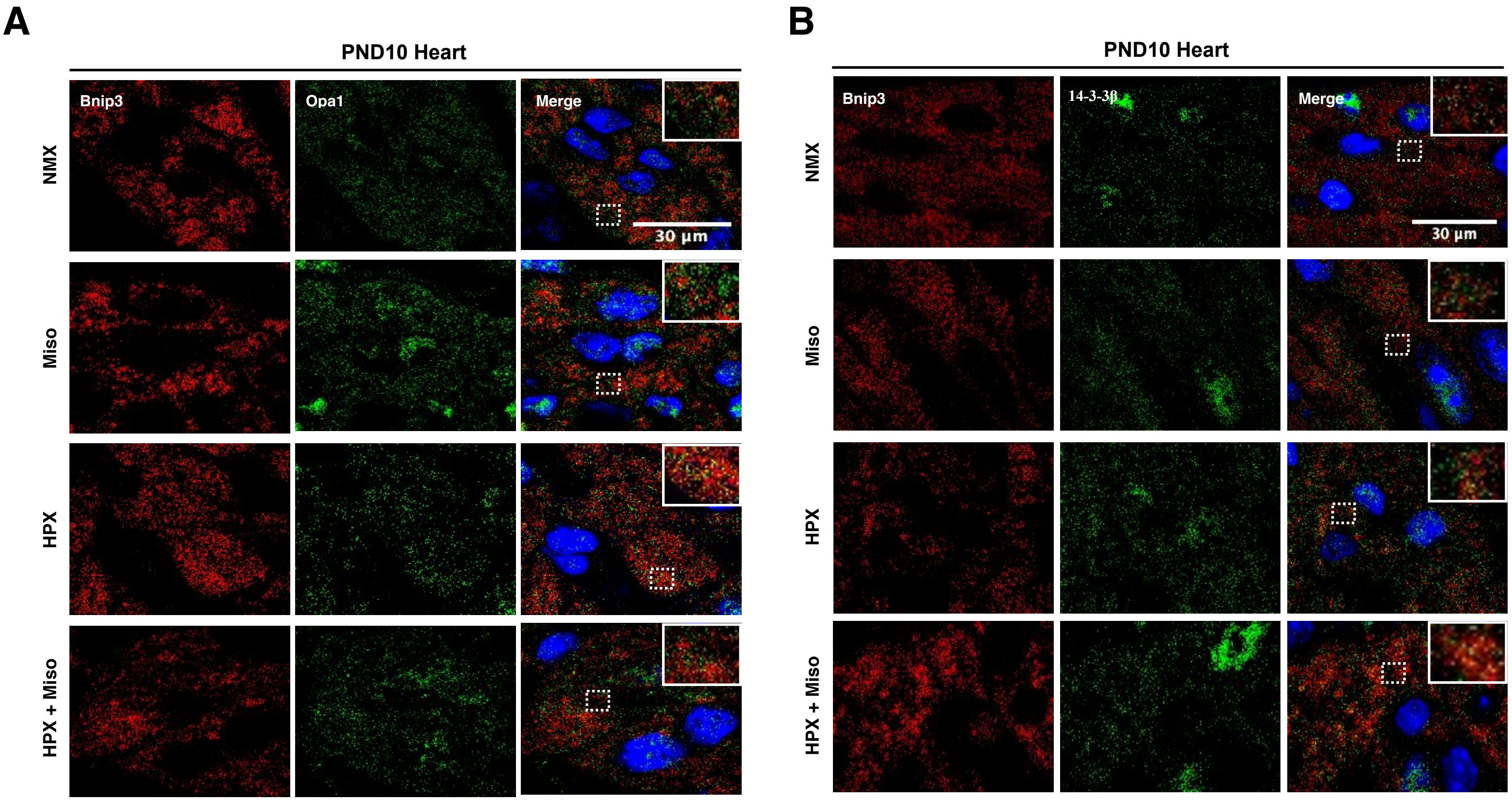


**Supplement 6. Misoprostol alters Bnip3's interactions with Opa1 and 14-3-3β. (A)** PND10 hearts from mice exposed to hypoxia (10% $O_2$) ± 10 µg/kg misoprostol from PND3-10, stained with DAPI (Blue) and probed for Bnip3 (Red), and Opa1 (green). Hearts were imaged via confocal microscopy. **(B)** PND10 hearts from mice treated as in (A) and stained with DAPI (Blue) and probed for Bnip3 (Red), and 14-3-3β (green). Hearts were imaged via confocal microscopy.
